# TxConformal: Controlling False Discoveries in AI-Driven Therapeutic Discovery

**DOI:** 10.64898/2026.04.27.721076

**Authors:** Ying Jin, Kexin Huang, Nathaniel Diamant, Kerry R. Buchholz, Steven T. Rutherford, Nicholas Skelton, Tommaso Biancalani, Gabriele Scalia, Jure Leskovec, Emmanuel J. Candès

## Abstract

Artificial Intelligence (AI) is transforming therapeutic discovery by scoring a large set of promising candidates and prioritizing a shortlist for further investigation. Quantifying the reliability of AI scores and preventing false positives among selected candidates is key to the efficiency of the discovery process. Conformal prediction (CP) has emerged as a popular tool for guiding such prioritization, especially via the conformal selection framework to control false discovery rates (FDR) in selecting top-ranked candidates under distributional shift^1, 2^. However, deploying these advances in real-world therapeutic discovery remains challenging: distribution shifts are difficult to quantify and correct in high-dimensional biomedical data, and practical workflows often require flexible error metrics. Here, we present TxConformal, a general framework for trustworthy decision making when building shortlists using AI scores. TxConformal adjusts for distribution shift by balancing the hidden representations in AI models and then provides confidence measures for true discoveries of target biological properties. These confidence measures, interpretable as p-values, can be used in conjunction with statistical multiple testing procedures to derive selection decisions with limited false positives or to estimate the errors in given selection decisions. TxConformal controls the false positive rate in six real-world tasks spanning various therapeutic discovery stages, modalities, and AI models with realistic data splits. When selecting promising combinatorial genetic perturbations, TxConformal nearly halves false-positive selections compared to baseline methods, substantially reducing unnecessary experimental costs by tens of thousands of dollars. When selecting stable protein structures under mutant shifts, TxConformal identifies about 10 times more proteins than baseline methods at stringent thresholds when running at a target FDR level of 10%, recovering over 90% of valuable candidates that baseline methods miss due to unaccounted distribution shifts. Furthermore, we demonstrate that TxConformal robustly supports various alternative error metrics suitable for resource-constrained settings. Finally, in a prospective fixed-budget virtual screening campaign for novel antibiotic discovery, TxConformal predicted false positives in close agreement with experimental outcomes, with substantial improvements over simple baselines.

## 1 Introduction

Artificial Intelligence (AI) is transforming drug discovery, driving advancements across diverse stages of therapeutic development such as target identification^3, 4^, virtual screening^5–7^, and clinical outcome prediction^8, 9^. AI systems trained on large-scale datasets can predict properties of biological entities with increasingly remarkable precision^10–12^. Given the significant costs of experimental validation of drug candidates and therapeutic hypotheses, it has become common to leverage predictions from AI models to prioritize small subsets of the most promising candidates for follow-up validation. This approach significantly reduces costs while increasing the chances of identifying high-potential candidates. The success of subsequent experimental cycles critically depends on the reliability of such AI-driven selection decisions: a falsely prioritized drug candidate in early stages can waste significant resources and time in later developments^1, 13, 14^ (Figure 1a). Therefore, reliable selection frameworks are essential for addressing critical questions in drug discovery, such as determining how many candidates to select to ensure a desired success rate or control the number of errors. This can enhance decision-making, shorten experimental cycles, and ultimately lower costs throughout the pipeline (Figure 1b).

**Figure 1:**
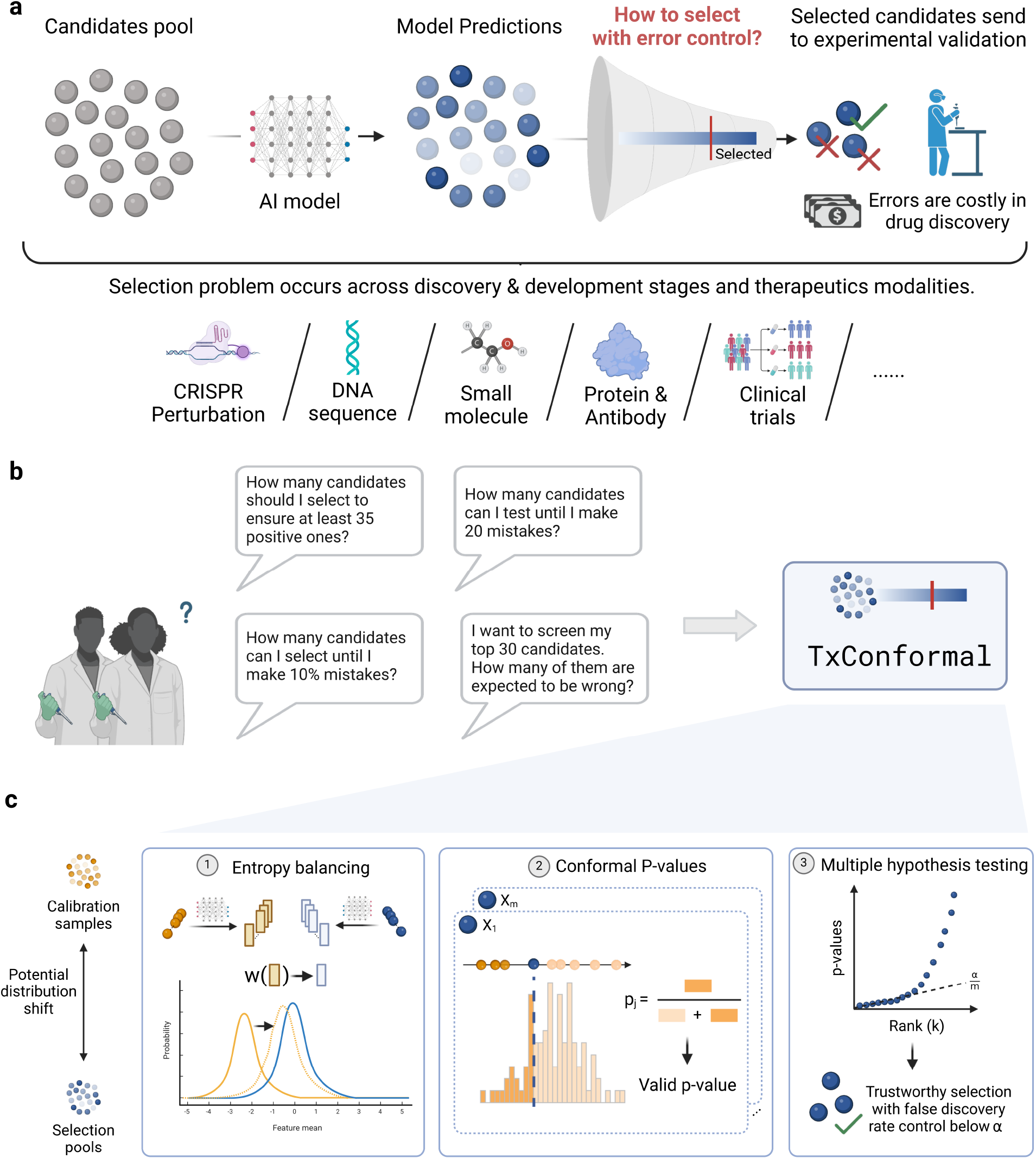
Overview of TxConformal for error-controlled candidate selection in drug discovery. **(a)** AI models predict properties of candidates to prioritize a subset for experimental validation. Errors at this stage can lead to significant waste of resources, highlighting the need for rigorous error control. This selection problem spans various drug discovery modalities, including CRISPR perturbation, DNA sequences, small molecules, proteins, and clinical trials. **(b)** TxConformal addresses key selection challenges, such as determining the number of candidates needed to ensure a desired success rate, estimating false positives in a selected subset, and setting tolerable error thresholds. **(c)** The framework consists of three components: (1) Entropy balancing, which adjusts for distribution shifts between calibration and selection pools; (2) Conformal p-values, enabling statistically valid error quantification; and (3) Multiple hypothesis testing, ensuring false discovery rate control. Together, these elements provide robust and reliable selection guarantees, improving decision-making in real-world drug discovery pipelines.

*Conformal prediction* has proven a popular framework for addressing such reliability issues. As an example, consider a molecular activity prediction task. Given the predicted activity of a new drug candidate, conformal prediction leverages existing drugs with experimentally evaluated activity scores to produce a range of values with a formal coverage guarantee: if the new drug is generated from the same distribution as the existing drugs, then with high probability, the experimental activity of the new drug falls within this range^15–17^. Such a range effectively quantifies the reliability of the AI prediction and has been used to suggest highly potent candidates^18, 19^. However, selecting AI-generated discoveries with controlled errors involves more than simply applying conformal prediction to individual drugs drawn from the same distribution^1, 14, 20^. We highlight two sources of distribution shift that complicate the quantification of errors when selecting promising candidates.

The first is the *selection effect*, where the set of selected candidates systematically differ from the broader candidate pool as they typically appear more promising. The core technical challenge here is not just ensuring the overall accuracy of the AI model—it is about providing rigorous statistical guarantees specifically for the *subset* of candidates chosen for experimental validation^14^. For example, selecting 100 molecules out of millions for lab testing using a model with 90% overall accuracy does not guarantee that 90 of them will actually pass experimental validation.

The second is the *distributional gap* between the known labeled data and the candidate pool where new discoveries must be made. Prioritization models are typically trained on examples of known effective candidates, then applied to predict outcomes on novel candidates with potentially very different characteristics^14^. In drug discovery, these shifts routinely occur, and they are critical to improving exploration and supporting discovery. For example, virtual screening seeks new scaffolds beyond the training set, which could be linked to novel mechanisms or improved activity and safety profiles^21–23^, while de novo molecular design methods are used to produce biomolecules outside natural domains through the optimization of specific properties such as binding or safety^24–27^.

Addressing both issues can reduce the risk of costly false positives and help capture valuable candidates that might otherwise be overlooked, making experimental validation significantly more efficient and reliable. Recent advances in statistical methodology, especially the conformal selection framework^1, 2, 28, 29^, have extended conformal prediction to address these issues in theory, enabling false discovery rate (FDR) control for selection under certain types of distribution shift. Yet, these methods assume the analyst has access to accurate models of distribution shifts, which remains challenging in real-world deployments. Indeed, approaches based on estimating the density ratio between two distributions can break down in applications due to the high dimensionality of biochemical data, leading to unstable or extreme estimated ratios. Moreover, distribution shifts may have several sources, such as changes in chemical space or design strategies, yet it remains unclear how to establish a unified approach that addresses them all. Furthermore, practical drug discovery applications may prioritize different error control metrics beyond FDR, such as those adapting to experimental budget constraints, which are not supported by existing approaches.

Here, we present TxConformal, a framework for reliable risk assessment in the selection of biological instances based on predictions from AI models. Notably, TxConformal does not depend on the intricacies of the AI model and can thus be applied in broad settings. TxConformal addresses two critical challenges: (1) controlling diverse error metrics in selection decisions, and (2) adapting to unknown distribution shifts. By leveraging entropy balancing to adjust for discrepancies in the latent embeddings of neural network models between labeled and unlabeled data, TxConformal effectively corrects for the unknown distribution shifts between novel hypotheses and existing ones. Such corrections are integrated into the theoretical framework of conformal selection^1, 2^ to compute statistical confidence measures, interpretable as p-values, to quantify the likelihood of false positives within certain selection sets. TxConformal then uses these p-values to enable diverse types of error control through multiple testing procedures, which leads to reliable selection decisions.

We validate TxConformal’s efficacy across multiple therapeutic discovery tasks, including both retrospective benchmarks and prospective screening. In six real-world therapeutic discovery tasks spanning various stages, modalities, and AI models, we create realistic data splits to emulate practical scenarios. For combinatorial genetic perturbations, TxConformal nearly halves false-positive selections compared to baseline methods (5 vs. 12 false positives at 25% FDR), substantially reducing unnecessary experimental costs by tens of thousands of dollars. For selecting protein structures under mutant shifts, TxConformal identifies more than 10 times as many promising stable proteins as baseline methods at stringent thresholds (633 vs. 58 selections at 10% FDR), recovering over 90% of valuable candidates that baseline methods miss due to unaccounted distribution shifts. In an ADMET prediction task, TxConformal further supports practical deployment questions for planning experiments given error/budget constraints, enabling resource-constrained decision-making. Finally, motivated by TxConformal’s retrospective performance, we evaluated its prospective utility in a novel screening campaign. We applied TxConformal in a fixed-budget phenotypic virtual screening campaign against *Acinetobacter baumannii* to identify novel antibacterial molecules, with subsequent wet-lab validation. Despite substantial distribution shift between the historical labeled data and the prospective screening pool, TxConformal accurately identified the number of false discoveries, highlighting its utility to support virtual screening campaigns and, more broadly, improve the efficiency of therapeutic discovery under realistic prospective settings. In summary, TxConformal provides a versatile framework for principled decision-making throughout the AI-driven drug discovery pipeline.

## 2 Results

### 2.1 Overview of TxConformal

Consider a therapeutic discovery task that aims to identify promising candidates with a desirable biological property (e.g., a high binding affinity to a disease target) from a pool of new instances. Here, a *true discovery* is an instance whose unknown property (e.g., binding affinity), measured by a numerical value, passes a user-specified threshold if tested via a physical experiment, and the goal is to reliably draw discoveries before experimental validation of the property. In such scenarios, we typically have access to labeled instances—referred to as *calibration data*—whose biological properties have already been experimentally observed in previous experiments. We also assume access to an AI model that predicts the unknown biological properties of the candidates, although these predictions carry inherent uncertainty. For simplicity, we consider the case where the model is trained independently of both the labeled calibration data and the new candidate pool; if the model must be trained using available labeled data, we treat the calibration data as a random subset of all labeled instances separate from the training dataset.

Building on the conformal inference framework^15^, TxConformal provides rigorous statistical guarantees for controlling prediction errors when selecting candidates with unknown biological properties from the pool. Specifically, TxConformal selects a subset of candidates with a guarantee that at least a pre-specified fraction (e.g., 90%) will truly exhibit the desired biological property (a true discovery). Additionally, TxConformal supports other critical decision-making tasks, including controlling the number of false discoveries (candidates selected but later found not to exhibit the desired property), ensuring a minimum number of true discoveries, and estimating the false discovery rate of a selected candidate set.

The entire workflow of TxConformal (Figure 1c) consists of three modules: (i) estimating the distribution shift between the labeled calibration data and the unlabeled candidate pool we aim to select from, (ii) constructing confidence measures (called conformal p-values) for each new instance, and (iii) combining the p-values to derive the selection set or quantify selection errors. Next, we introduce each component, with details in the Methods.

#### Entropy balancing

TxConformal includes a component to account for the distribution shift between the calibration samples and test data, incorporating the representations learned by AI models into the entropy balancing algorithm^30^ in causal inference. Entropy balancing finds a set of weights that balance the expected value of statistics of the calibration and test data, such that relevant features appear similar across the two sets after reweighting the calibration data. TxConformal is based on the intuition that the learned representations from the deep learning models capture essential semantic information of the biological instances that allow not only predicting the outcome, but also explaining distributional differences across datasets. In TxConformal, the features include the predicted values, indicators for quantile-based bins, and embeddings from the deep learning prediction models. The learned weights are the maximum-entropy weights that match the empirical distribution of predicted values and sample mean of embeddings across the calibration and test data.

#### Conformal p-values

After obtaining the entropy balancing weights, TxConformal measures the statistical confidence in large responses, i.e., the desired property of the test samples, based on model predictions. For each test data, it computes a conformal p-value which quantifies how likely the unknown test label may surpass a user-specified threshold. At a high level, this is achieved by comparing the predicted value of the test sample against the predictions for the labeled calibration data using any function referred to as a conformity score^15^. While being built upon AI models, these p-values are valid in a familiar sense^1, 2^: the probability of having a small (undesirable) test label while observing a p-value smaller than any *α* ∈ (0, 1) is upper bounded by *α* under mild conditions (Methods). As with classical p-values, a small conformal p-value indicates a strong evidence that the unobserved test label is large, thereby grounding the discoveries with solid statistical evidence.

#### Multiple testing

Finally, TxConformal derives trustworthy discoveries by passing the conformal p-values onward to multiple testing procedures to produce a selection set (a list of discoveries) controlling meaningful error metrics. In our primary use case, we apply the Benjamini-Hochberg procedure^31^ to our p-values to determine a selection set that controls the false discovery rate (FDR, the expected proportion of false positives among the selected instances) below a nominal level. Controlling the FDR implies a guaranteed expected hit rate for the selected instances, such that follow-up validation efforts are mostly devoted to truly rewarding candidates. Within this process, we select test samples whose p-values are no greater than a threshold determined by TxConformal. In addition, TxConformal can use the conformal p-values to estimate false positives in a given selection set, determine how many discoveries are needed to meet a minimum number of true positives, and produce a list of discoveries with a limited number of false positives.

#### Theoretical guarantees

We establish theoretical analysis showing that TxConformal asymptotically achieves the desired guarantee, given that either the entropy balancing method accurately captures the distribution shift between calibration and test data, or the prediction model accurately predicts the property (Methods) (more specifically, if the hidden representations of the AI model capture either the density ratio or the property distribution). Although these conditions cannot be exactly verified in practice, our empirical results in real drug discovery tasks corroborate TxConformal’s robust error control.

### 2.2 TxConformal accurately controls the FDR in diverse selection tasks

We demonstrate the efficacy and robustness of TxConformal in five diverse therapeutic discovery tasks for identifying interesting instances with false discovery rate (FDR) control. The FDR is the expected fraction of false discoveries (i.e., selected instances whose unobserved labels are not sufficiently large) among the selected instances. Notably, each task presents a distinct type of realistic distribution shift between the labeled calibration data and unlabeled test data. In each task, we employ a state-of-the-art model for predicting the biological property of interest and apply Tx-Conformal to derive the selection set by constructing p-values with estimated distribution shift weights. The performance of TxConformal is compared with that of deriving the selection set based on conformal prediction without accounting for the selection effect or distribution shift (Methods, Section A.6). The baseline which does not account for selection effect represents the most common approach in applying conformal prediction to drug discovery^18, 19^.

Across all tasks, TxConformal consistently maintains tight FDR control across the entire spectrum of nominal levels. This robustness means that practitioners can select a nominal level appropriate for their application with confidence that the error control will hold. In contrast, the baseline method either underestimates the selection error leading to exceedingly high error rates, or overestimates the selection error leading to insufficient selection power (the ability to select test instances with the desired properties).

#### Selection of protein structures with desirable properties under mutant shift

Our first task focuses on selecting proteins with high stability, a crucial objective in protein engineering and therapeutic development. Protein stability plays a key role in ensuring the effectiveness of biologics, such as drugs that need to remain intact until delivered to their target, and is critical for industrial applications and enzyme design^32, 33^. Beyond drug development, designing stable proteins helps maximize yield in costly protein engineering experiments, especially by refining top candidates identified through broader initial screening^34^. Computational approaches produce predictions of protein stability with the sequence data as input, which are invaluable for prioritizing candidates when experimental resources are limited^35, 36^. However, robust selection methods are needed to ensure predictions lead to actionable insights.

This task introduces a challenging mutant shift, where the labeled calibration sets consist of proteins from four rounds of experimental design, while the test set includes top candidate proteins with single mutations (Hamming distance-1 neighbors)^37^. This setup challenges a model’s ability to generalize from a broad sampling of protein sequences and to localize this knowledge to select top candidates.

Across all nominal FDR levels, TxConformal reliably maintains the empirical FDR below the target level (Figure 2a). In contrast, the baseline methods, which do not account for selection effects (Conformal Baseline) or the mutant shift (TxConformal-Unweighted), either significantly underestimates or overestimates it. For example, at the nominal level of 0.1, TxConformal is able to confidently select 633 proteins (averaged over all replica), where on average 90% of the selected are true positives, while the baseline method (without accounting for selection effects) is only able to select 57.8 proteins on average, missing over 90% of promising candidates that could have been confidently identified (Figure 2c). In addition, only accounting for the selection bias without adjusting for distribution shifts (TxConformal-Unweighted) tends to overestimate the empirical FDR – such a method yields a realized FDR of 0.758 at a nominal level of 0.3, whereas TxConformal yields an FDR of 0.234. These results demonstrate TxConformal ‘s robustness in prioritizing stable proteins, making it a valuable tool for protein engineering under realistic experimental conditions.

**Figure 2:**
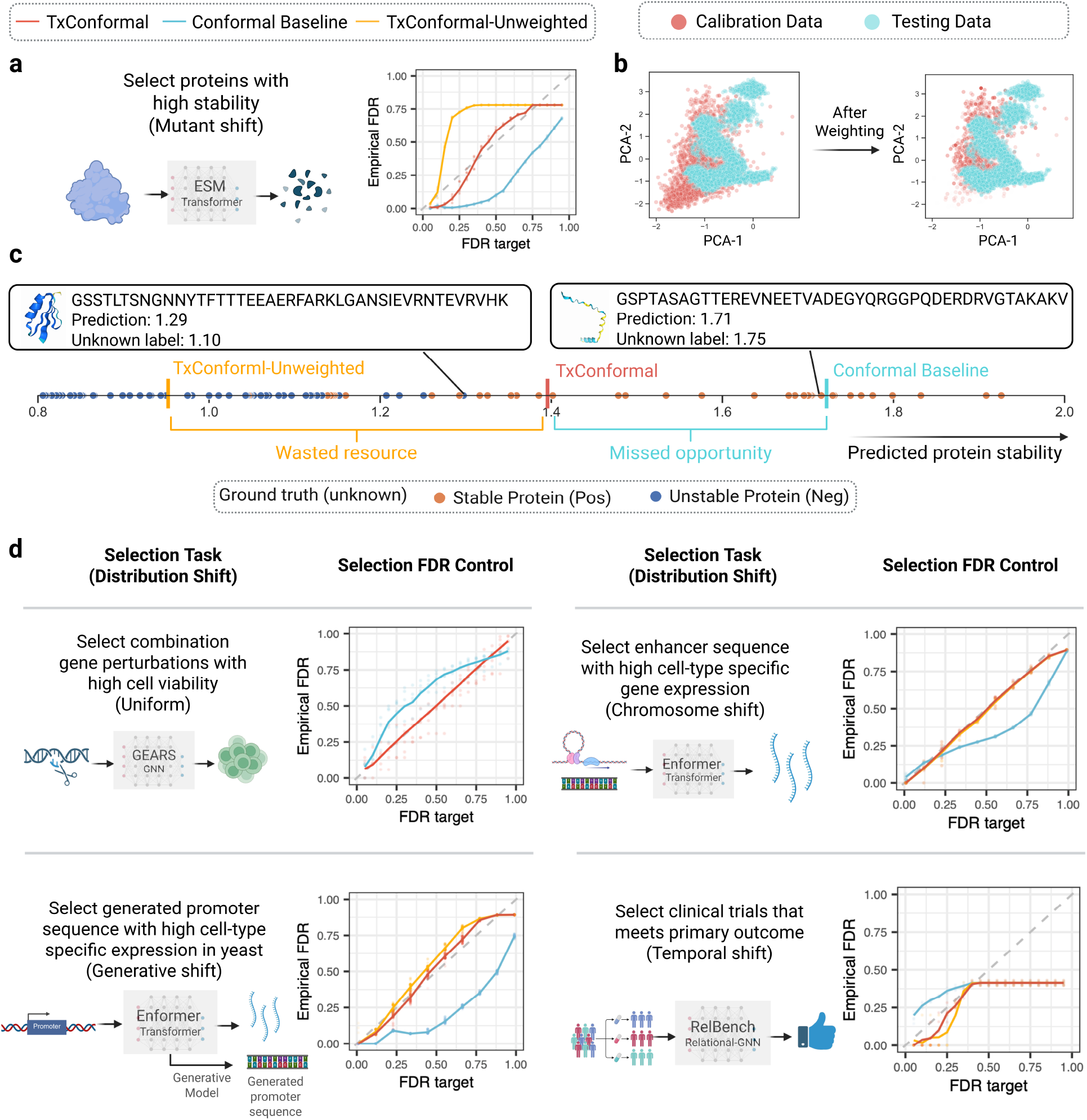
TxConformal accurately controls false discovery rate (FDR) across diverse therapeutic discovery tasks under distribution shifts. This figure illustrates five tasks and their corresponding FDR control performance. **(a)** The task of selecting proteins with high stability under a mutant shift, with TxConformal leveraging ESM Transformer to tightly control the empirical FDR below the target (red), while the conformal baseline (blue) and unweighted version (orange) fail to do so. **(b)** The first two principal components of calibration proteins and test (unlabeled) proteins before (left) and after (right) reweighting. **(c)** Selection decisions in one run of TxConformal in the protein selection task; each dot is the predicted stability of a test protein, where a brown dot is a true discovery and a blue dot is a false one. TxConformal finds a selection threshold at round 1.4, while the conformal baseline misses opportunities due to conservativeness, and the unweighted version wastes follow-up resources by including too many false positives. **(d)** Performance of TxConformal in four other selection tasks: selection of combination gene perturbations with high cell viability under uniform sampling (no distribution shift) using the GEARS GNN model; selection of AI-generated promoter sequences with high cell-type-specific expression under a generative shift, guided by Enformer predictions; selection of enhancer sequences with high cell-type-specific gene expression under a chromosome-based shift, using predictions from Enformer; selection of clinical trials meeting primary outcomes under a temporal shift, with predictions from RelBench. Across all tasks, TxConformal demonstrates superior robustness and accuracy in FDR control under diverse real-world distribution shifts, significantly outperforming the baseline methods.

As an illustration, consider the protein sequence “GSSTLTSNGNNYTFTTTEEAERFARKL-GANSIEVRNTEVRVHK,” where the AI model predicts a stability score of 1.29, while the ground truth is 1.1. This sequence is not highly stable and should therefore be excluded from the short-list to avoid wasting experimental resources. The TxConformal-Unweighted method incorrectly includes it, whereas TxConformal correctly excludes it. Conversely, for the sequence “GSPTASAGTTEREVNEETVADEGYQRGGPQDERDRVGTAKAKV,” which has a high stability label of 1.75 and a model prediction of 1.71, the Conformal Baseline fails to include it in the shortlist, representing a missed opportunity. In contrast, TxConformal successfully selects it. These examples illustrate how the tighter uncertainty calibration of TxConformal achieves an improved balance between avoiding wasted resources and capturing valuable candidates.

#### Selection of combinatorial genetic perturbations

Genetic interactions between small sets of genes underlie complex cellular phenotypes, driving processes critical to precision cell engineering, such as tumor suppression and cell differentiation^38^. Identifying gene-gene interactions can enhance the design of combination therapies and improve the success rate of clinical trials^39, 40^. While experimental techniques have become more efficient at sampling perturbation outcomes, the combinatorial explosion of potential gene combinations makes computational approaches indispensable. Machine learning models have emerged as powerful tools for predicting perturbation effects. These models take the list of perturbed genes as input and predict the post-perturbed cellular response, enabling researchers to prioritize combinations for experimental validation^41, 42^. However, these predictions are prone to errors, necessitating robust selection methods to identify promising gene combinations for wet-lab validations.

The upper left panel of Figure 2d demonstrates this task. We use a uniform split between calibration and test data, reflecting the early-stage exploration of genetic perturbations where combinations are sampled broadly without prior knowledge of their biological significance. This setup mirrors practical scenarios where researchers seek to systematically uncover novel interactions and emergent properties by exploring a diverse range of perturbations. We employ the GEARS model, a state-of-the-art graph neural network designed to predict the effects of gene combinations on cell viability^41^. Using these predictions, TxConformal does not need distribution shift adjustment since the gene combinations are randomly assigned to the calibration and test sets. Across all nominal FDR levels, TxConformal reliably controls the empirical FDR below the nominal level. In contrast, the baseline method, which does not account for selection effects, mostly underestimates the FDR, leading to excessive false positives. For example, at nominal level *α* = 0.1, the baseline method has a false positive rate of 20%, meaning that the fraction of incorrectly selected perturbations doubles what is sought. The tight error control of TxConformal implies a reduction by tens of thousands of dollars of experimental costs. These results highlight the value of Tx-Conformal in exploring gene combinations by prioritizing configurations aligned with model predictions.

#### Selection of enhancer sequence under chromosome shift

Our third task focuses on selecting enhancer sequences that elicit high cell-type-specific gene expression, a critical step toward understanding gene regulation and its role in cellular identity. Enhancers are regulatory DNA elements that drive gene expression in a context-specific manner^43^. Identifying active enhancers can reveal mechanisms underlying tissue-specific functions, developmental processes, and disease states^44^, with profound implications for designing gene therapies. While experimental assays like STARR-seq or ChIP-seq can profile enhancer activity^45^, the scale of the enhancer landscape makes computational prediction crucial for prioritization. Machine learning models such as Enformer have shown promise in predicting enhancer activity from DNA sequence data^46–48^. Since predictions contain errors, reliable selection of promising enhancers worthy of follow-up studies is crucial.

In the upper right panel of Figure 2d, we employ a chromosome-based split between calibration and test data in this task to reflect practical scenarios where enhancers are identified in regions of the genome not covered by the training data^46^. Such shifts are common in enhancer studies due to the diversity of regulatory regions across chromosomes. TxConformal adjusts for distribution shifts between chromosome splits based on the embeddings in the Enformer model^49^, and then uses the predictions from the model to construct selection sets. Across all nominal FDR levels, TxConformal consistently controls the empirical FDR below the target threshold. In contrast, the baseline method, which does not account for the selection effects, overestimates it, especially at large nominal levels, reducing selection power. For example, with a target FDR of 0.45, TxConformal declares 1228 enhancers as confident positive discoveries on average, while the baseline method only declares 976 enhancers as positive discoveries, missing over 20% of opportunities. These results demonstrate TxConformal ‘s effectiveness in prioritizing enhancer sequences under realistic experimental conditions, offering robust and precise guarantees for downstream validation.

#### Selection of AI-generated promoter sequences

The fourth task focuses on selecting generated promoter sequences with high cell-type-specific expression, a key problem in synthetic biology and gene regulation engineering (lower left panel of Figure 2d). Promoters are regulatory DNA sequences that control gene expression^50^, and designing promoters with desired regulatory activity enables constructing synthetic gene circuits^51^, which would have great potential in therapeutic applications that require context specificity^46^. Recent advances in generative AI models have made it possible to computationally design novel promoters, sampling from the vast combinatorial space of candidate sequences^52^. However, their effectiveness in accurately achieving the desired biological function must be validated experimentally. Given computationally designed regulatory sequences, robust selection methods are needed to prioritize those to be experimentally tested, avoiding wasted resources while retaining promising candidates.

This task introduces a *generative shift*, where the calibration set includes natural promoter sequences, and the test set contains synthetic promoters generated by a generative model^52^. This setup reflects real-world scenarios in synthetic biology, where models must generalize from natural data to evaluate and select novel, artificially designed sequences. Using predictions from Enformer^49^, TxConformal adapts to the generative shift based on the embeddings from the same model and constructs the selection sets. Across all nominal FDR levels, TxConformal consistently controls the empirical FDR below the target threshold. In contrast, the baseline method, which does not account for the selection effects or the generative shift, largely overestimates the empirical FDR and misses many confident instances. The effect of neglecting the distribution shift is visible in this case, since the Conformal-Unweighted method, which only adjusts for the selection effect, violates FDR control. These results highlight the robustness of TxConformal in prioritizing high-performing synthetic regulatory elements, offering significant utility in therapeutic and biotechnology applications.

#### Selection of clinical trials with preferable outcomes under temporal shift

The fifth task concerns selecting clinical trials that would meet their primary outcome, a crucial step in evaluating trial success and optimizing resource allocation in clinical research. Predicting the success of clinical trials is a key application in health data analytics, with direct implications for time and cost efficiency in drug development^9, 53^. In this task, machine learning models predict the likelihood of success based on the information about the trial before it happens. This task exemplifies predictive modeling in clinical outcomes, where reliable selection is essential to mitigate prediction errors and optimize decision-making.

As demonstrated in the lower right panel of Figure 2d, this task naturally introduces a temporal shift, where the calibration data comprises earlier clinical trials, while the test set includes more recent (or future) trials, reflecting real-world scenarios where models must generalize to newer data^54^. RelBench, a relational graph neural network designed for analyzing structured trial relational databases, provides predictions for trial success based on trial characteristics in the database^55^. Using RelBench predictions and embeddings, TxConformal constructs selection sets and adapts to the temporal shift between calibration and test data. Across all nominal FDR levels, TxConformal consistently maintains the empirical FDR below the target level. In contrast, the baseline method, which does not account for selection effects or temporal shifts, underestimates FDR—resulting in excessive false positives. For instance, when TxConformal controls the FDR below the nominal level of 10%, the baseline method incurs an FDR of over 25%. In addition, the TxConformal-Unweighted method, which only addresses the selection effect, misses opportunities for identifying promising future trials. These results demonstrate the effectiveness of TxConformal in prioritizing promising clinical trials under realistic, time-evolving conditions.

### 2.3 TxConformal informs false positives in diverse deployment scenarios

We further demonstrate the wide applicability of TxConformal in offering flexible error control guarantees in various deployment scenarios when coupled with diverse multiple testing strategies, using a sixth task on selecting candidate compounds with desirable activity. While the challenges of selection and distribution shift are similar, TxConformal achieves types of guarantees distinct from the FDR control.

We focus here on the task of selecting compounds with high CYP2D6 inhibition rates, an example of ADMET (Absorption, Distribution, Metabolism, Excretion, and Toxicity) property prediction, which is critical during lead optimization in drug discovery^56, 57^. CYP2D6 is a key enzyme involved in drug metabolism, and compounds with high inhibition rates can lead to adverse drug-drug interactions or toxic buildup^58^. Identifying such compounds early allows drug developers to filter them out, improving the safety and efficacy of therapeutic candidates. In doing so, it is equally important to avoid mistakenly discarding safe compounds, as the cost of missed opportunities is especially high at this stage. With molecular property prediction models facilitating in silico profiling of ADMET properties by screening large compound libraries^59^, robust selection frameworks are critical to ensure the reliable prioritization of candidates using these models. We consider scenarios with a scaffold shift, where the calibration set consists of compounds with diverse chemical scaffolds, while the test set focuses on novel scaffolds not seen during training^60^. This setup reflects a realistic challenge in drug discovery, where generalization to unseen chemical spaces is necessary for identifying new leads. Our experiments use embeddings from AttentiveFP^61, 62^ to train the prediction model on a holdout set of data and learn the distribution shift adjustment between data splits.

#### Scenario 1: Controlling the false discovery rate

The first scenario has the same goal, namely, FDR control while adapting to the distribution shift introduced by the scaffold split (Figure 3, row 3). When the scientist is interested in achieving an FDR below 0.1 (i.e., at most 10% in the selection set fail to meet the desired property), TxConformal informs them that 62 top-ranked instances can be confidently selected (“Result” column). TxConformal reliably maintains the empirical FDR below the target level across all nominal levels (“Validity” column), providing reliable decision support at any desired error control level that scientists may find appropriate. The baseline method (which does not account for scaffold shifts and selection effects) underestimates the selection errors and yields excessive false positives; we do not depict it here for consistent visualization across all panels. For instance, at the nominal FDR level of 10%, the baseline method incurs an FDR over 20%, which doubles the target error budget. These results further highlight TxConformal’s robustness in supporting the selection of safe and effective compounds under realistic experimental conditions.

**Figure 3:**
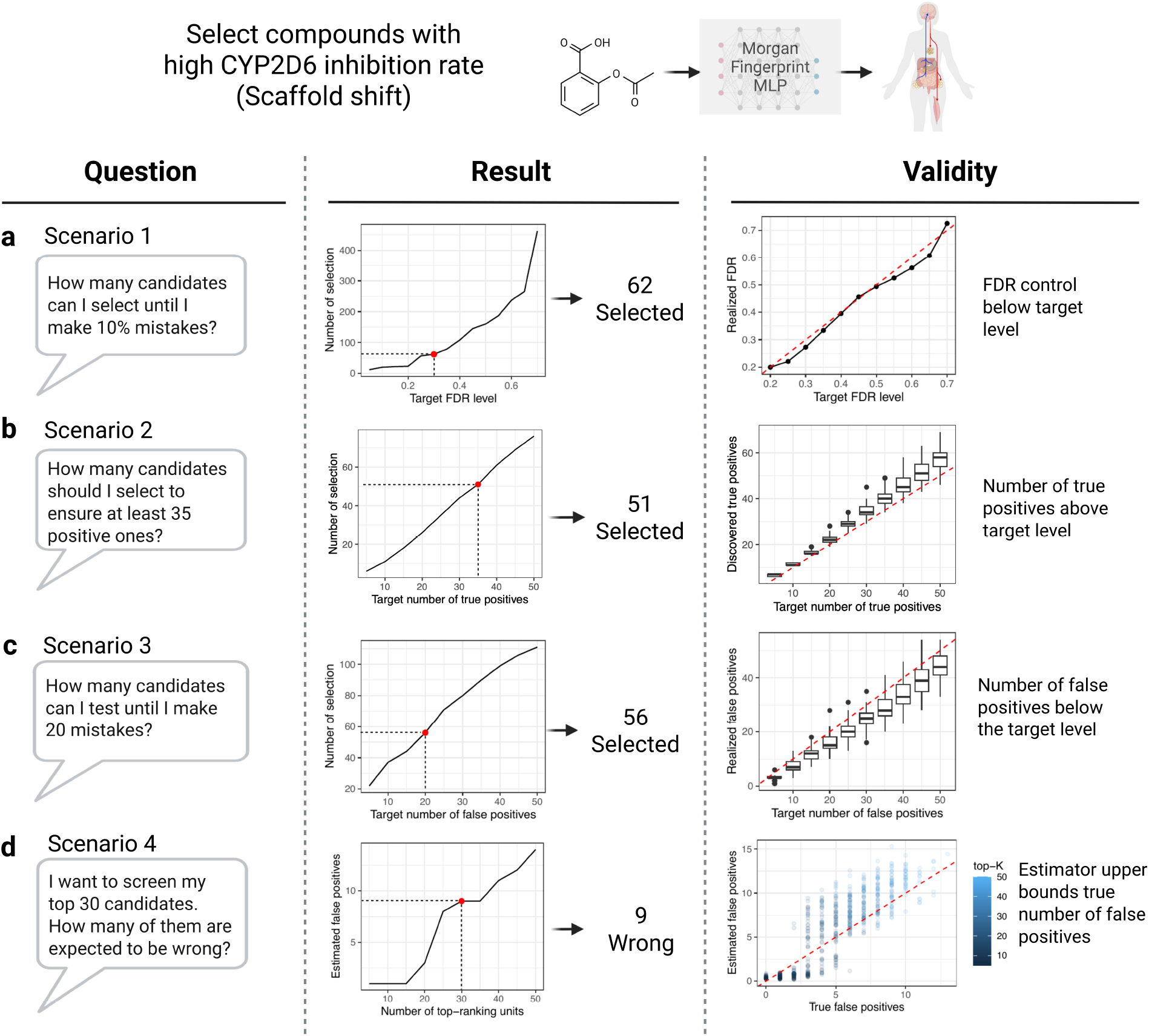
TxConformal achieves error control guarantees in diverse deployment scenarios. This figure demonstrates how TxConformal addresses distinct error control guarantees across multiple deployment scenarios using conformal p-values in the task of selecting compounds with high CYP2D6 inhibition rates, where a label of one indicates a true discovery. **(a)**: Controlling the false discovery rate (FDR). The Result plot shows the required number of selections to control FDR at various levels (x-axis: target FDR level, y-axis: number of selections), with 62 candidates selected at a target FDR level of 30%. The Validity plot demonstrates that realized FDR is consistently below the target. **(b)**: Ensuring a minimum number of true positives. The Result plot shows the required number of selections to achieve a given number of true positives with high probability (x-axis: target number of true positives, y-axis: number of selections), with 51 candidates selected to ensure at least 35 true positives on average. The Validity plot validates that the number of discovered true positives meets or exceeds the target in most cases. **(c)**: Controlling the expected number of false positives. The Result plot illustrates the required number of selections to maintain false positives below a specified target (x-axis: target number of false positives, y-axis: number of selections), with 56 candidates selected to ensure no more than 20 mistakes on average. The Validity plot confirms that the realized false positives are typically below the target. **(d)**: Estimating false positives in a given selection set. The Result plot depicts the expected false positives in a top-ranked subset (x-axis: number of top-ranking units, y-axis: estimated false positives), with 9 false positives estimated in a set of 30 candidates. The Validity plot validates that estimated false positives serve as an upper bound for true false positives in most cases. These results highlight the versatility and reliability of TxConformal in addressing a range of error control needs across different scientific workflows.

#### Scenario 2: Determining the number of selections needed for a minimum of true positives

The second type of guarantee we address is the true positive count, defined as the expected number of true positives in the selection set. This task is critical in drug discovery to ensure that sufficiently many confirmed candidates are available for further exploration. Thresholding an estimated number of true positives based on the conformal p-values, TxConformal produces a selection set with a guarantee on the minimum number of true discoveries. In this way, TxConformal quantifies the number of candidates that must be examined to guarantee a specified minimum of true discoveries. As a concrete example, when a scientist is interested in obtaining at least 35 correct selections, TxConformal estimates that 51 instances need to be screened to ensure this goal (“Result” column). Across multiple independent runs of the procedure, and as we vary the target number of true positives, the realized number of true positives in the TxConformal selection set lies above the target number in most cases (“Validity” column). In other words, TxConformal offers a likely lower bound for the number of candidates needed to screen so as to have at least 35 correct selections.

#### Scenario 3: Controlling the expected number of false positives

The third type of guarantee TxConformal addresses is the false positive count, defined as the expected number of false positives in the selection set. Using the conformal p-values to estimate and cap the number of false positives, TxConformal produces a selection set with bounded expected number of false positives. When the scientist has a budget of how many false positives they can afford—for example, in settings where validation is expensive and time-consuming—TxConformal informs them of the extent to which they can explore the candidate list within the error budget. For example, when the scientist is interested in how many instances can be selected while expecting fewer than 20 mistakes, TxConformal estimates that at most 56 drug candidates can be tested (“Result” column). Across multiple runs of our procedure, the number of false positives lies below the target values most of the time, showing the ability of TxConformal in informing trustable selections in resource-constrained settings (“Validity” column). While our theoretical framework (Methods) establishes the bounds on the expected number of false positives, we indeed observe high-probability validity in our experiments, with the number of false positives bounded below the target level in most replications. In this example, TxConformal offers a likely upper bound for the number of candidates that can be screened before making at least 20 mistakes.

#### Scenario 4: Estimating false positives in a given selection set

In our final scenario, we consider a setting where a shortlist of candidates has been constructed by taking the first *K* instances ranked by predicted value, which reflects a standard procedure in many applications. Instead of suggesting a shortlist, the use of TxConformal here is to estimate—and provide a likely upper bound for—the number (or fraction) of false positives in the selection set. This post-hoc process offers scientists a quantification of how many erroneous hits to expect in the shortlist. In one run of this procedure, TxConformal estimates that there are 9 false positives in the shortlisted set of top 30 candidates (“Results” column). Across multiple runs of the procedure, the estimated number of false positives from TxConformal indeed serves as a likely upper bound for the unknown number of false positives in most cases (“Validity” column).

### 2.4 TxConformal accurately predicts false positives in prospective deployment

Prospective deployment is the setting in which trustworthy error quantification is most critical. Under a fixed experimental budget, compounds must be prioritized for synthesis and assay before any outcomes are observed, and each false positive directly consumes experimental resources. We therefore evaluated TxConformal in a prospective virtual screening to identify antibacterial compounds (Fig. 4a).

**Figure 4:**
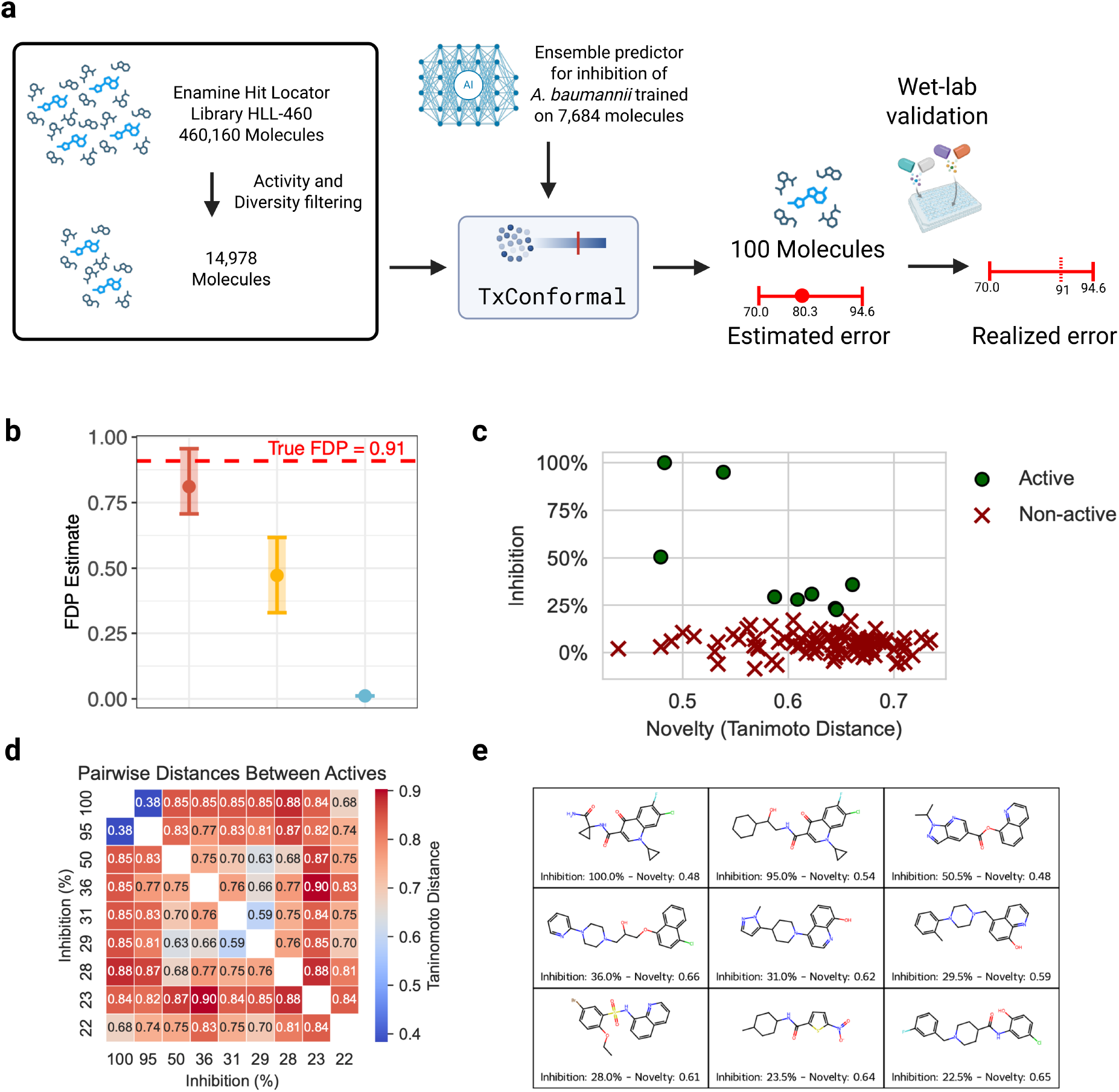
TxConformal accurately predicts false positives in prospective deployment. **(a)**: Overview of the prospective virtual-screening workflow. Starting from 460,160 molecules in the Enamine Hit Locator Library (HLL-460), activity-, novelty- and diversity-based filtering yielded 14,978 candidate molecules. TxConformal was then applied using an ensemble graph neural network-based predictor of *A. baumannii* inhibition growth trained on a public library. Candidate molecules were ranked by predicted score, and the top-100 compounds were prioritized for synthesis and experimental validation. Before experimental validation, Tx-Conformal estimated 80.3 false positives among the selected molecules (90% confidence interval [70.0, 94.6]), compared with 91 false positives realized experimentally. **(b)**: Estimated false discovery proportion (FDP) for the selected set from TxConformal, its unweighted variant and a proxy implied by the conformal baseline, compared with the realized FDP of 0.91 from experimental validation. Error bars indicate 90% confidence intervals where available. **(c)**: Experimental inhibition versus molecular novelty, quantified as Tanimoto distance to the training and calibration molecules. Active compounds were defined as having mean inhibition ≥ 20% across two replicates; green circles indicate actives and red crosses indicate non-actives. **(d)**: Pairwise Tanimoto distances among the nine validated active molecules, showing that the diversity-filtering step preserved substantial structural diversity within the hits. **(e)**: Chemical structures of the nine validated active molecules, annotated with measured inhibition and novelty (minimal Tanimoto distance from any active moledules in the labeled data).

This task is particularly important in antibiotic discovery, where multidrug-resistant bacteria create an urgent need for new therapeutics^63^. Recent work has shown that deep-learning-guided virtual screening can uncover structurally novel antibacterial hits from ultralarge chemical libraries^22, 64, 65^. Yet the extremely low hit rates in this setting, together with the scale of the candidate space and the small number of compounds that can be advanced to experimental validation, make prospective assessment of shortlisted candidates essential.

We focus on prioritizing molecules with antibacterial activity against *Acinetobacter baumannii*, a Gram-negative pathogen designated a highest-priority target for antibiotic development owing to the urgent need for new treatment options^66^.

The central practical question in this setting is not only which molecules to prioritize, but whether the expected number of false positives in a shortlisted set can be estimated before wet-lab validation. In the following, we demonstrated how TxConformal can support this phase in a prospective virtual screening setting.

For this experiment, a graph neural network-based model was trained to predict antibacterial activity given a molecular structure, following recent works^22, 64^. In particular, we used an ensemble of two GIN-based models (Methods Section A.8) which exhibit robust FDR control under various synthetic shifts in the training process (SI Figure 5). TxConformal also uses their predictions and learned representations to estimate reweighting factors to reduce the discrepancy in predicted activity between the labeled and unlabeled datasets (lower panel of SI Figure 6). The model was trained on public high-throughput *Acinetobacter baumannii* data^65^, and the same dataset was used for subsequent calibration. The unlabeled candidate pool comprised an activity-, novelty-, and diversity-filtered subset of the Enamine Hit Locator Library (HLL-460) (Methods Section A.8).

**Figure 5.**
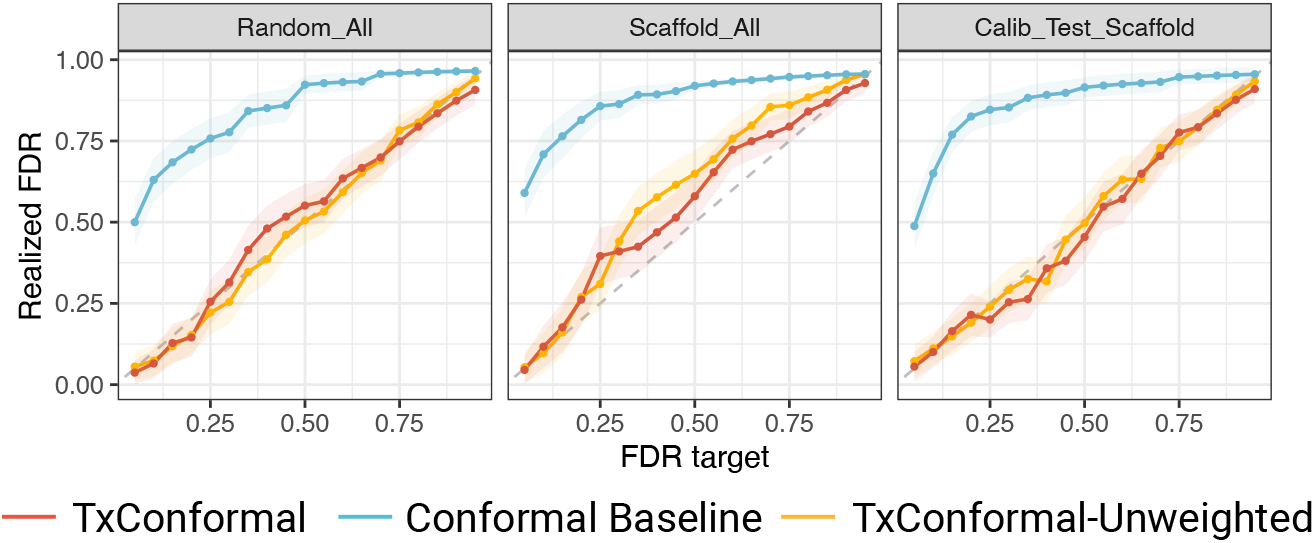
Empirical FDR on the test fold using the NCB data over *N* = 100 splits, with the predictions and hidden embeddings from the ensemble models trained on the training fold. Left: results when the training, calibration (labeled) data and test (unlabeled) data are split randomly. Middle: results when the training, calibration, and test folds are all with distinct scaffolds. Right: results when the training data is a random subsample from the entire dataset, and calibration and test folds are split by scaffolds.

**Figure 6.**
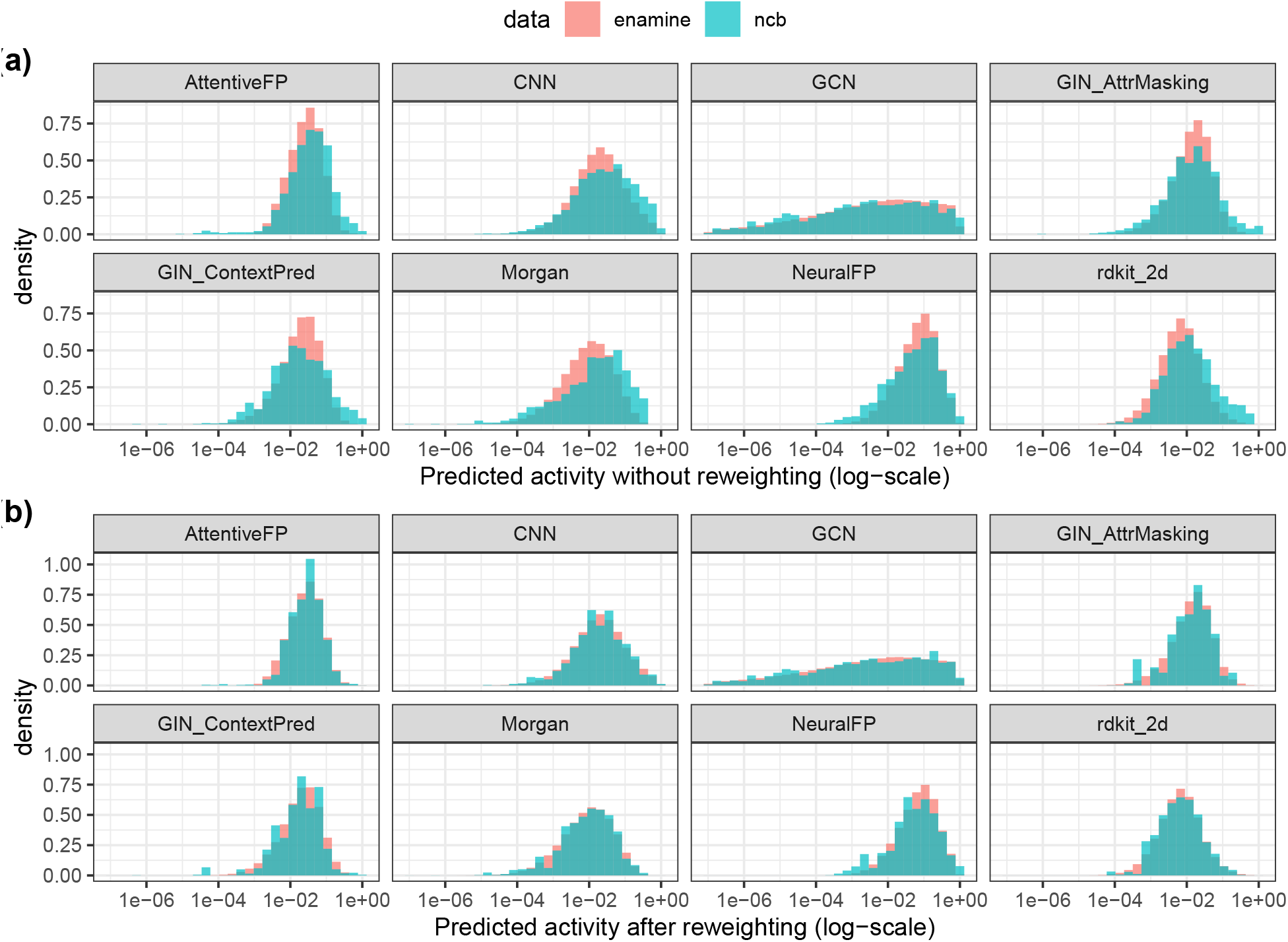
TxConformal produces empirical weights that balance predicted activities for all the prediction models. (a) Histograms of model predictions in the labeled data (ncb, blue) and prospective new samples where discoveries are to be made (enamine, red). Each panel corresponds to the predicted activities from one of the eight candidate models. (b) Histograms after weighting using weights produced by TxConformal using the selected ensemble models.

This deployment scenario induces substantial distribution shift between labeled and unlabeled molecules and is representative of virtual-screening-based hit discovery aimed at identifying novel chemical scaffolds^22^. In particular, the Enamine candidates are chemically distinct from the three source libraries underlying the training dataset, and the final screening pool is further shaped by multi-step filtering. As a result, and consistently with previous work^22^, several property prediction models show pronounced shifts in predicted activity between the labeled and unlabeled datasets (upper panel of SI Figure 6). This makes the task a stringent test of whether TxConformal can remain well calibrated under realistic deployment shift and provide actionable estimates of false positives before synthesis and experimental testing.

We apply TxConformal to produce conformal p-values that quantify the statistical confidence in high antibacterial activity against *Acinetobacter baumannii* for each candidate compound, where the distribution shift between the labeled data and unlabeled data is adjusted based on the ensemble model representations. Based on the experimental budget of *K* = 100 compounds, we select the top-100 compounds with the smallest conformal p-values (strongest statistical confidence) for synthesis. We then apply TxConformal as in Scenario 4 in Figure 3 to estimate the number of false positives in a given selection set.

Prior to experimental validation, TxConformal produces a point estimate of 80.29 false positives, with the 90% confidence interval based on normal approximation of [70.00, 94.60]. In contrast, the unweighted baseline (without distribution shift adjustment) produces a point estimate of 46.80 false positives, with a 90% confidence interval of [32.50, 61.11]. The conformal prediction baseline widely used in the literature^18, 67^, which makes selection decisions based on conformal prediction sets of the unknown property at a fixed marginal coverage level, is not directly applicable here, because it does not naturally estimate the number of false positives. In practice, applying this strategy, a scientist would use prediction sets with marginal coverage 1 − *α* = 0.9889 to decide on pursuing these top-100 compounds, resulting in a overly optimistic assessment of the error rates.

Selected compounds were experimentally validated to investigate their activity, measuring inhibition growth against *Acinetobacter baumannii*. Experimental validation identified nine active molecules, defined as compounds with mean inhibition ≥ 20% across two replicates, yielding a realized FDP of 0.91. Notably, this realized error fell within the interval estimated by TxConformal, but far outside that of the unweighted variant, highlighting the importance of explicitly accounting for distribution shift in prospective deployment (Fig. 4b). Although the most active compounds tended to be less novel, which is consistent with a common trade-off in virtual screening^64^, several validated hits remained substantially novel relative to the training and calibration molecules (Tanimoto distance ≥ 0.6 from any calibration actives; Fig. 4c). The diversity-filtering step also preserved substantial structural variation among the actives, as reflected by generally large pairwise Tanimoto distances and the distinct chemotypes recovered among the nine validated hits (Fig. 4d,e). Together, these results show that TxConformal can provide practically useful estimates of false positives in a realistic fixed-budget screening campaign under challenging distribution shifts, well-aligned with experimental outcomes.

## 3 Discussion

We introduced TxConformal—a general framework for trustable candidate selection in therapeutic discovery tasks that bridges the gap between predictive accuracy and actionable decision-making. By integrating entropy balancing with conformal inference and multiple testing procedures, TxConformal offers rigorous error control, even under challenging scenarios such as distribution shifts and selection biases. Our comprehensive evaluation across six diverse therapeutic discovery tasks and additional deployment scenarios demonstrates that TxConformal not only tightens false discovery rate (FDR) control but also adapts flexibly to alternative error metrics, providing critical support for resource-constrained experimental pipelines. Our framework is able to quantify uncertainty in a manner that is directly interpretable as a statistical significance measure. The construction of conformal p-values, in tandem with adaptive weighting via entropy balancing, enables rigorous control of the proportion of false positives in a selection set.

Despite these promising results, several potential limitations warrant consideration. First, the performance of TxConformal is inherently tied to the quality of the underlying AI models and their learned latent representations. The entropy balancing step assumes that these representations capture key structural features of the biological instances; if the representations fail to accurately predict the distribution shift or the unknown labels, the adjustment for distribution shifts may be incomplete. From this point of view, continued advances in foundation models across biochemical modalities, driven by pretraining on large-scale data^68, 69^, are expected to enhance the quality of these representations. Moreover, the framework relies on the availability and representativeness of calibration data—a prerequisite that may not always hold, especially in settings characterized by limited or biased experimental observations. From this perspective, iterative experimental strategies that incrementally explore the candidate space^70–72^ could help overcome this limitation. Third, the computational overhead of entropy balancing and multiple testing, though manageable in our experiments, could scale poorly with extremely large test sets. Finally, TxConformal requires predefined thresholds for “desirable” biological properties, which may not align with dynamic discovery goals. Methods that can detect and protect against insufficiency in these aspects would be valuable for further enhancing our framework.

Looking ahead, several avenues of investigation could extend the impact of TxConformal. One promising direction is to evaluate its performance in more diverse prospective studies and real-world clinical settings, where the stakes of decision-making are highest. Additionally, exploring alternative error metrics—such as family-wise error rate control or tailored metrics for specific therapeutic contexts—could further enhance the framework’s versatility. There is also potential to adapt TxConformal to other domains beyond drug discovery, including genomics, synthetic biology, and even non-biomedical applications like materials science, where selection decisions based on AI predictions are equally critical.

## Data availability

The datasets used in the analysis are publicly available online. The dataset in the high-stability protein selection task is available in https://torchdrug.ai/docs/api/datasets.html#stability. The dataset in the gene perturbation selection task is available in https://www.ncbi.nlm.nih.gov/geo/query/acc.cgi?acc=GSE133344. The dataset in the enhancer sequence selection task is available in Supplementary Table 2 from https://pmc.ncbi.nlm.nih.gov/articles/PMC11525185/ with download link https://pmc.ncbi.nlm.nih.gov/articles/instance/11525185/bin/41586_2024_8070_MOESM4_ESM.txt. The dataset in the promoter sequence selection task is available at https://zenodo.org/records/4436477. The dataset in the clinical trial selection task is available in https://relbench.stanford.edu/datasets/rel-trial/. The dataset in the experiments in Figure 3 is available in https://tdcommons.ai/single_pred_tasks/adme/#cyp-p450-2d6-inhibition-veith-et-al. The molecule library in the prospective deployment (Enamine Hit Locator Library, HLL-460) is from https://enamine.net/compound-libraries/diversity-libraries/hit-locator-library-460, whose predicted activities and hidden embeddings are shared in the project repository for reproduction.

## Code availability

The Python implementation of TxConformal and the scripts for reproducng the analysis in this work, along with usage examples, are available at https://github.com/ying531/TxConformal.

## Acknowledgements

E.C. was supported by the Office of Naval Research grant N00014-24-1-2305, the National Science Foundation grant DMS-2032014, and the Simons Foundation under award 814641. J.L. gratefully acknowledge the support of NSF under Nos. OAC-1835598 (CINES), CCF-1918940 (Expeditions), DMS-2327709 (IHBEM), IIS-2403318 (III); Stanford Data Applications Initiative, Wu Tsai Neurosciences Institute, Stanford Institute for Human-Centered AI, Chan Zuckerberg Initiative, Amazon, Genentech, GSK, Hitachi, SAP, and UCB. K.H. was supported by Stanford Bio-X fellowship.

## Authors contribution

Y.J. and K.H. designed and developed the framework. Y.J. developed the methods and the theoretical results. Y.J, K.H., and N.D. implemented the analysis. All authors discussed the results and contributed to the final manuscript.

## A Methods

The Methods section is organized as follows. Sections A.1 to A.5 describe the general framework of TxConformal, including (1) an overview of notation, (2) the construction of entropy balancing weights, (3) conformal p-values, (4) selection procedures producing Figure 2 and Figure 3. Section A.6 describes the baseline methods under comparison. Section A.7 describes the datasets and models used in the analysis, together with data splitting methods to create the realistic distribution shifts and evaluation strategies.

### A.1 Notations and setup

We assume access to a set of identically and independently distributed (i.i.d.) labeled data 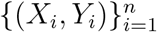 (referred to as calibration data) and a set of i.i.d. unlabeled data 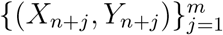 (referred to as test data) with 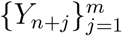 unobserved. Each sample *i* is a biological instance (such as a molecule, a protein, a clinical trial), *X*_*i*_ ∈ 𝒳 is the feature vector used for making the prediction (such as the chemical structure of a molecule, the sequence of amino acids of a protein, and features of a clinical trial), and *Y*_*i*_ ∈ ℝ is a numerical value describing the property of interest. Our method applies to any prediction model *f*: 𝒳 → ℝ which takes features *X* as input and produces a prediction *f* (*X*) for the unknown property *Y* . The only requirement is that the training process of *f* is independent of the calibration and test data; for instance, it is pre-trained on an independent set of data and does not utilize any data in the calibration and test sets. Throughout our applications, we choose *f* as state-of-the-art deep learning models.

To model the distribution shift between labeled and unlabeled data, we assume 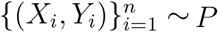 for some unknown distribution *P* while 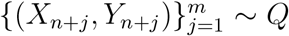 for another unknown distribution *Q*. We impose the covariate shift assumption: d*Q/*d*P* (*x, y*) = *w*^∗^(*x*) for some unknown function *w*^∗^: 𝒳 → ℝ^+^. This means the features capture all the distributional difference between labeled and unlabeled data. Put differently, the distribution of the response given the features is invariant across the two datasets, which is suitable for many biological problems. As an example, in virtual screening, the binding affinity *Y* conditional on the chemical structure *X* does not depend on whether the sample is from the calibration or test set.

#### Assumption 1

*The test data are i*.*i*.*d. from an unknown distribution* 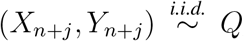 *and the calibration data are i*.*i*.*d. from another unknown distribution* 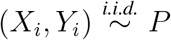. *The true density ratio between the test and calibration data exists, and is only a function of the features; that is, w*^∗^(*x*) = d*Q*(*x, y*)*/* d*P* (*x, y*).

We focus on tasks where large values of the unknown labels 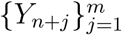 are of interest. Given user-specified thresholds 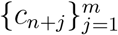, a test point with *Y*_*n*+*j*_ *> c*_*n*+*j*_ is considered a true discovery, which is unknown since *Y*_*n*+*j*_ is not yet evaluated. We study various error control guarantees in producing a selection set 𝒮 or estimating the quality of a given selection set.

### A.2 Entropy balancing weights

The first step of TxConformal is to estimate the covariate shift function *w*(·). Tailored to the nature of biological discovery tasks and designed to harness the representation power of AI prediction models, TxConformal offers a stable and efficient framework that can be applied across a wide range of problems.

While standard approaches in the literature rely on density ratio estimation^73–75^ (that is, separately estimating *P* (*x*) and *Q*(*x*) and taking the ratio), estimating densities can be challenging for high-dimensional, unstructured features. Therefore, TxConformal directly estimates the function *w*(*x*) using insights from observational causal inference^30^. Specifically, we learn the weights *w*_*i*_ for *i* = 1, …, *n* which balance the distribution or moments of certain key features *ϕ*(*X*) ∈ ℝ^*p*^ between the two groups. The features *ϕ*(*X*) should be predictive of both the response and the likelihood of being a labeled or unlabeled data. Based on this idea, TxConformal combines the predicted values, deciles of the predicted values (among both unlabeled and labeled data), and hidden embeddings in the last layer of the deep learning model as the initial transformed features. Next, principal component analysis (PCA) is used to remove redundant linear components in the initial features; we then obtain the final features *ϕ*(*X*) after standardizing each feature to a unit standard deviation in the calibration data. The learned weight vector ***w*** = (*w*_1_, …, *w*_*n*_) is the solution to the following optimization problem:

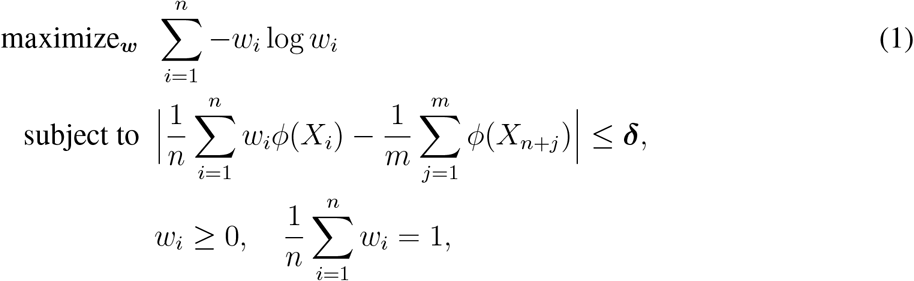

where ***δ*** = (*δ*_1_, …, *δ*_*p*_) is a vector of nonnegative tolerance levels for the “imbalance” between two groups after reweighting, and for any vector ***z*** ∈ ℝ^*p*^, |***z***| is the vector of magnitudes. When ***δ*** = 0, the solution to the above optimization program takes the form 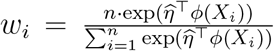 for some 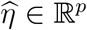. We thus set the learned weight function as 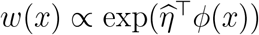 . In our experiments, we keep ***δ*** = 0, which makes the optimization program feasible most of the time. In cases where it is not, we gradually discard the principal components explaining less than 5% of the variance until the optimization program becomes feasible. If the program is still infeasible after discarding all such principal components, we gradually increase the tolerance levels to *δ*_*j*_ ≡ *δ* ∈ {0.001, 0.005, 0.01} until the feasible set of the optimization program is non-empty.

### A.3 Construction of conformal p-values

Recall that a test point with *Y*_*n*+*j*_ *> c*_*n*+*j*_ is considered a true positive. TxConformal quantifies the confidence in such a large outcome via conformal p-values^1, 2, 15^.

We wrap the prediction model with a monotonically increasing score function *V*: 𝒳 × ℝ → ℝ obeying *V* (*x, y*) ≤ *V* (*x, y*^′^) for any *x* ∈ 𝒳 and *y, y*^′^ ∈ ℝ, *y* ≤ *y*^′^. In addition, assume access to a function *w*(·) where *w*(*x*) estimates the density ratio *w*^∗^(*x*). Then, for each test point *j* ∈ {1, …, *m*}, we define a p-value following Jin and Candès (2023)^2^:

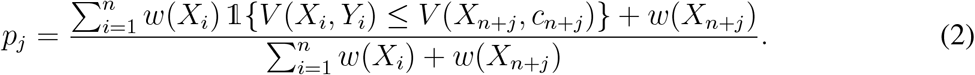

For instance, one may set *V* (*x, y*) = *y* − *f* (*x*) for continuously-valued or binary outcomes. The intuition of the p-value is as follows: if the weights *w*(*X*_*i*_) consistently estimate the true density ratio *w*^∗^(*x*) and we substitute *V* (*X*_*n*+*j*_, *c*_*n*+*j*_) with *V* (*X*_*n*+*j*_, *Y*_*n*+*j*_), then the p-value would be dominated by a uniform distribution between [0, 1]. Thus, observing an extremely small p-value indicates that *V* (*X*_*n*+*j*_, *c*_*n*+*j*_) is smaller than what *V* (*X*_*n*+*j*_, *Y*_*n*+*j*_) would typically be, which further implies evidence for *Y*_*n*+*j*_ *> c*_*n*+*j*_ since *V* is non-decreasing in the second argument.

Throughout this paper, for classification problems with *Y* ∈ {0, 1}, we take *V* (*x, y*) = *y* · *M* − *f* (*x*) for *M* = 100 ^1, 2^. For regression problems with *Y* ∈ ℝ and a constant threshold *c*_*n*+*j*_ ≡ *c* ∈ ℝ, we take *V* (*x, y*) = *M* 𝟙{*y > c*} + *c* 𝟙{*y* ≤ *c*} − *f* (*x*) with *M* = 100. While we primarily use a constant threshold in this study, our framework also accommodates settings with non-constant *c*_*n*+*j*_’s. In such cases, we assume access to calibration thresholds {*c*_*i*_} and construct calibration scores *V* (*X*_*i*_, *Y*_*i*_, *c*_*i*_) and test scores *V* (*X*_*n*+*j*_, *c*_*n*+*j*_, *c*_*n*+*j*_), where *V* (*x, y*; *c*) = *M* 𝟙{*y > c*} + *c* 𝟙{*y* ≤ *c*} − *f* (*x*) for a sufficiently large constant *M >* 0.

Our conformal p-values are similar to classical statistical p-values in the sense that ℙ(*p*_*j*_ ≤ *t, Y*_*n*+*j*_ ≤ *c*_*n*+*j*_) ≤ *t* holds approximately for *t* ∈ (0, 1) given reliable estimation of the density ratio or the prediction model (see Section C in the supplementary material). Throughout our theory, we place the following basic assumptions on the data generating process.

#### Assumption 2

*We assume w*^∗^(*X*_*i*_) *and w*(*X*_*i*_) *are bounded by a constant almost surely. The features ϕ*(*x*) *in the entropy balancing optimization program with δ* = 0. *The features ϕ*(·) *and the scores V* (·) *are obtained independently of the calibration and test data*.

Since *w*(*x*) is subject to estimation error, we need to resort to asymptotic analysis to study the performance of our procedure. The following theorem establishes the robust behavior of individual conformal p-values with two routes to statistical validity.

#### Theorem 3.

*When P* = *Q and we run* TxConformal *with w*(*x*) ≡ 1, *it holds in finite sample that* ℙ(*p*_1_ ≤ *t, Y*_*n*+1_ ≤ *c*_*n*+1_) ≤ 1 *for any monotone score function. With general covariate shift (Assumption 1), when w*^∗^(·) = *w*(·), *then* ℙ(*p*_1_ ≤ *t, Y*_*n*+1_ ≤ *c*_*n*+1_) ≤ 1 *for any monotone score function. Otherwise, under Assumption 1, upon setting δ* = 0 *in the entropy balancing algorithm* (1), *it holds that* 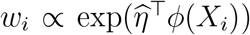 *for some* 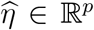. *Also, there exists some fixed* 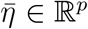 *such that* 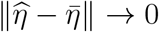 *as n, m* → ∞. *Under Assumptions 1 and 2, for any t* ∈ [0, 1], *the limit of* ℙ(*p*_1_ ≤ *t, Y*_*n*+1_ ≤ *c*_*n*+1_) *as n, m* → ∞ *exists, and* lim_*n,m*→∞_ ℙ(*p*_1_ ≤ *t, Y*_*n*+1_ ≤ *c*_*n*+1_) ≤ *t under any of the following conditions:*

1. *The weight function is log-linear in ϕ*(*x*), *i*.*e*., *w*^∗^(*x*) ∝ exp(*η*^⊤^*ϕ*(*x*)) *for some η* ∈ ℝ^*p*^.
2. *The score function is conditionally invariant, i*.*e*., *there exists an unknown, fixed function V* ^∗^ *such that* 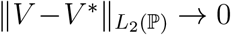 *as n, m* → ∞ *and the distribution of V* ^∗^(*X*_*i*_, *Y*_*i*_; *c*_*i*_) *conditional on X*_*i*_ *is invariant across i* = 1, …, *n* + 1 *almost surely*.

Intuitively, Theorem 3 reveals that the p-value is “valid” when either the weight function can be recovered by the entropy balancing program (which is the case under condition (i)), or the score function based on the prediction model converges to a function with invariant conditional distribution (which is the case if *V* ^∗^ captures the conditional c.d.f. of *Y*_*n*+*j*_). In the first case, the distribution shift is effectively adjusted by entropy balancing, whereas in the second, the distribution shift has no effect if our model is perfect for the conditional distribution of the outcomes.

In the next steps, we will use these p-values as if they were classical p-values to draw discoveries using hypothesis testing techniques and show the guarantees of such procedures.

### A.4 Controlling the FDR via Benjamini-Hochberg procedure

We now address the problem of selecting a subset 𝒮 ⊆ {1, …, *m*} with false discovery rate (FDR) control. The procedure in this part corresponds to Figure 2.

The FDR is defined as the expected proportion of falsely discovered instances among the selection set:

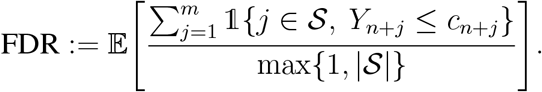

Given a pre-specified nominal level *α* ∈ (0, 1), TxConformal produces a selection set 𝒮 such that its FDR is below *α*. The set 𝒮 is the shortlist of promising candidates TxConformal identified for subsequent experimental validation. By definition, controlling the FDR ensures a prescribed proportion of these follow-up experimental validations are devoted to truly desired instances in expectation.

To achieve FDR control, TxConformal applies the Benjamini-Hochberg procedure^2, 31^ to the conformal p-values 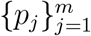 constructed in the preceding step. Specifically, we define

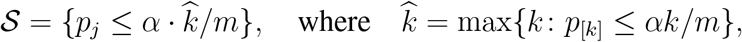

where *p*_[1]_ ≤ *p*_[2]_ ≤ · · · ≤ *p*_[*m*]_ are the order statistics of the p-values 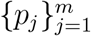 and *α* is the target FDR.

#### Theorem 4.

*When P* = *Q and we run* TxConformal *with w*(*x*) ≡ 1, *it holds in finite sample that* FDR ≤ *α for any monotone score function. Under Assumption 1, we have* lim_*n,m*→∞_ FDR:= lim_*n,m*→∞_ 𝔼[|𝒮 ∪ ℋ_0_|*/*(1 ∨ |𝒮|)] ≤ *α with* ℋ_0_:= {*j* ∈ [*m*]: *Y*_*n*+*j*_ ≤ *c*_*n*+*j*_} *under any of the following three conditions: conditions (i), (ii) in Theorem 3, and*

*(iii) There exists some fixed function V* ^∗^ *such that* 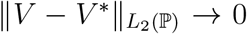. *In addition, the conditional c*.*d*.*f. of the score function at* 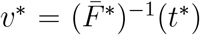 *is linear in ϕ*(*x*), *i*.*e*., ℙ(*V* ^∗^(*X*_*i*_, *Y*_*i*_) ≤ *v*^∗^ | *X*_*i*_ = *x*) = *β*^⊤^*ϕ*(*x*) *for some constant β* ∈ ℝ^*p*^ *for all i* ∈ [*n* + *m*]. *Here, we define t*^∗^ = sup{*t* ∈ [0, 1]: 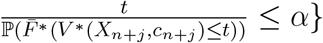 *and* 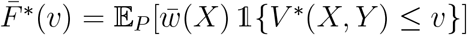, *and* 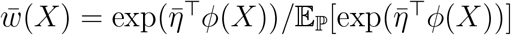 *for* 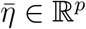 *in Theorem 3*.

Intuitively, these conditions show that the FDR can be controlled in different scenarios: (i) either the features capture the distributional differences between the calibration and test data, (ii) or they capture the distribution of the score, (iii) or the prediction model (score) accurately estimates the conditional distribution of *Y*_*i*_ given *X*_*i*_. These conditions motivate us to build the features based on the hidden representations in the deep learning prediction models. Note that condition (ii) concerns the construction of the scores based on the conditional distribution of *Y*_*i*_. In the binary case (*Y* ∈ {0, 1} and *c* = 0), it reduces to *f* (*x*) being consistent for ℙ(*Y* = 1 | *X* = *x*). For regression problems, in practice, it is still most convenient to use a learned point prediction instead of learning the conditional c.d.f. function, which has also demonstrated reliable performance.

### A.5 Controlling other error notions

Coupling the conformal p-values with other multiple testing procedures, TxConformal controls other notions of errors (Figure 3). In this part, we detail these procedures and the technical conditions for achieving the desired guarantees. Our theory establishes similar consistency conditions as those in Theorem 4, which also motivate building the features based on the learned representations in TxConformal.

#### Scenario 2: Determining the number of selections needed for a minimum of true positives

In this scenario, the goal is to ensure a minimum number of expected true positives. Given a target number *K* ∈ ℕ^+^, TxConformal produces a selection set 𝒮 ⊆ {1, …, *m*} such that approximately,

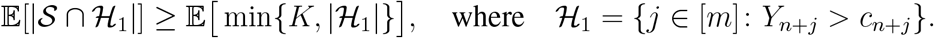

In the right-hand side, we take the minimum of the target number and the actual number of positive instances for tractability.

TxConformal achieves this goal by estimating from p-values the number of false positives and true positives. Specifically, we define

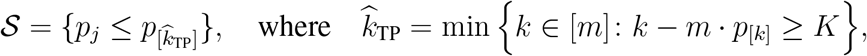

and min(∅) = *m*. Intuitively, *m* · *p*_[*k*]_ is an upward-biased estimate of how many false positives we have among the *k* test instances with the smallest p-values, and therefore *k* − *m* · *p*_[*k*]_ is a downward-biased estimate of the number of true positives in the most promising *k* test instances (with smallest p-values).

##### Theorem 5.

*Consider the asymptotic scheme n, m* → ∞ *and K/m* → *ρ for some constant ρ >* 0. *Then* lim inf_*n,m*→∞_ |𝒮 ∩ ℋ_1_|*/* min{*K*, |ℋ_1_|} ≥ 1 *in probability under any of the three conditions: conditions (i), (ii) in Theorem 3, and*

*(iii) There exists a fixed function V* ^∗^ *such that* 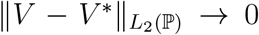. *In addition, the conditional c*.*d*.*f. of the score function at* 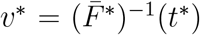 *is linear in ϕ*(*x*), *i*.*e*., ℙ(*V* ^∗^(*X*_*i*_, *Y*_*i*_) ≤ *v*^∗^ | *X*_*i*_ = *x*) = *β*^⊤^*ϕ*(*x*) *for some constant β* ∈ ℝ^*p*^ *for all i* ∈ [*n* + *m*]. *Here, we define* 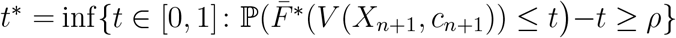 *where* 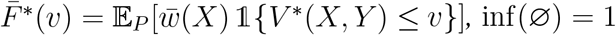 *and* 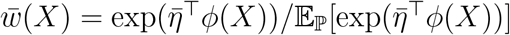 *for* 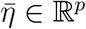 *in Theorem 3*.

#### Scenario 3: Controlling the expected number of false positives

In this scenario, the goal is to produce a selection set with an upper bound on the expected number of false positives. Concretely, given a target number *K* ∈ ℕ^+^, TxConformal produces a selection set 𝒮 ⊆ {1, …, *m*} such that approximately,

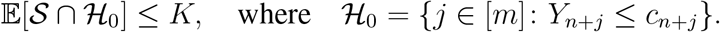

TxConformal achieves so by similarly estimating the number of false positives among the most promising instances with smallest p-values. Specifically, we define

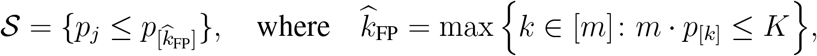

and max(∅) = 0. Similar to Scenario 2, *m* · *p*_[*k*]_ is a conservative estimate of the number of false positives in the most promising *k* test instances. Capping this estimate at the target number *K* yields our selection set.

##### Theorem 6.

*Consider the asymptotic regime m, n* → ∞ *and K/m* → *ρ for some constant ρ >* 0. *Then* lim sup_*n,m*→∞_ |𝒮 ∩ ℋ_0_|*/K* ≤ 1 *in probability under any of the following three conditions: conditions (i), (ii) in Theorem 3, and*

*(iii) Suppose there exists some fixed function V* ^∗^ *such that* 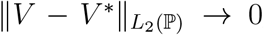, *and define* 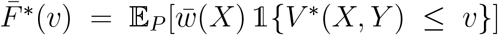. *The conditional c*.*d*.*f. of the score function at* 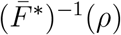 *is linear in ϕ*(*x*), *i*.*e*.,, *and* ℙ(*V* ^∗^(*X*_*i*_, *Y*_*i*_) ≤ *v*^∗^ | *X*_*i*_ = *x*) = *β*^⊤^*ϕ*(*x*) *for some constant β* ∈ ℝ^*p*^ *for all i* ∈ [*n* + *m*].

#### Scenario 4: Estimating false positives in a given selection set

Finally, we address scenarios with a fixed budget for one batch of *K* experimental validations, and have decided to select the *K* test instances with the highest prediction values from the model *f* . Since we rank with prediction values, we assume here *c*_*n*+*j*_ ≡ *c* for some constant *c >* 0; otherwise the ranking shall naturally depend on *Y*_*n*+*j*_ − *c*_*n*+*j*_.

In this scenario, TxConformal leverages p-values using a specific conformity score function to produce a conservative estimate of the number or fraction of false positives in the top-*K* selection list. We use the score *V* (*x, y*; *c*) = *M* 𝟙{*y > c*} + *c* 𝟙{*y* ≤ *c*} − *f* (*x*), so that test units with highest prediction values *f* (*x*) have the smallest conformity score *V* (*x, c*) = −*f* (*x*). We then define the conformal p-values as in (2). The quantity of interest is the number of false positives 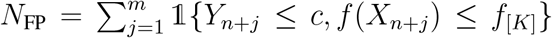 (where *f*_[*K*]_ is the *K*-th order statistic of 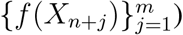. Our estimate for *N*_FP_ is given by 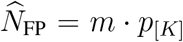, where *p*_[*K*]_ is the conformal p-value for the least promising test point in the selection set. Similarly, the false discovery proportion (FDP) in the list, FDP:= *N*_FP_*/K*, can be estimated via 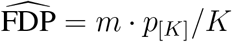.

##### Theorem 7.

*Consider the asymptotic regime n, m* → ∞ *and K/m* → *ρ for some constant ρ >* 0. *Then* 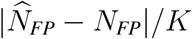 *converges to zero in probability under any of the three conditions:*

i. *The weight function is log-linear in ϕ*(*x*), *i*.*e*., *w*^∗^(*x, y*) ∝ exp(*η*^⊤^*ϕ*(*x*)) *for some η* ∈ ℝ^*p*^.
ii. *The conditional distribution of the prediction on the null is invariant, i*.*e*., *for any v*, ℙ(*Y*_*i*_ ≤ *c, f* (*X*_*i*_) ≤ *v* | *X*_*i*_ = *x*) *is invariant across i* ∈ [*n* + *m*] *and x* ∈ 𝒳 .
iii. *The conditional c*.*d*.*f. of the prediction at its ρ-th quantile, denoted as q*_*ρ*_, *is linear in ϕ*(*x*), *i*.*e*., ℙ(*f* (*X*_*i*_) ≤ *q*_*ρ*_ | *X*_*i*_ = *x*) = *β*^⊤^*ϕ*(*x*) *for some constant β* ∈ ℝ^*p*^ *for all i* ∈ [*n* + *m*].

### A.6 Baseline methods under comparison

Finally, we detail the two baseline methods under comparison in Section 2.2 (Figure 2). The Conformal Baseline method (referred to as CB hereafter) does not address distribution shift or selection effect, and TxConformal-Unweighted method (referred to as TU hereafter) ignores distribution shift.

CB follows existing literature on conformal prediction for drug discovery, deriving selection decisions directly from marginally valid conformal prediction sets^18, 67, 76^. Given the calibration data 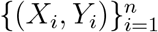, test samples 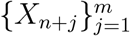, and target error rate *α* ∈ (0, 1), a one-sided conformal prediction set^15^ is first built for each test sample:

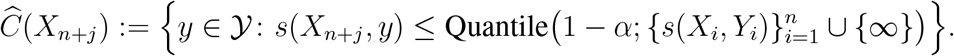

Here the outcome domain is 𝒴 = {0, 1} for classification tasks, and 𝒴 = ℝ for regression tasks, and Quantile(1 − *α*; 𝒱) denotes the empirical (1 − *α*)-th quantile of 𝒱 for any finite set 𝒱. In addition, the conformity score *s*: 𝒳 → 𝒴 is any function that measures the “conformity” of an outcome value *y* to a model prediction^15^. To construct a one-sided conformal prediction set of the form 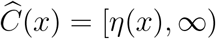, ∞) which provides evidence for large outcomes, we use commonly discussed approaches. For classification tasks where 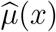 is an estimate of ℙ(*Y* = 1 | *X* = *x*), we set 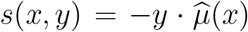 for regression tasks where 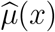 estimates 𝔼[*Y* | *X* = *x*], we set 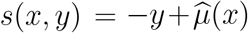. We then select all test samples *X*_*n*+*j*_ whose prediction set lies above the threshold *c*_*n*+*j*_,i.e., 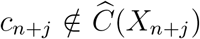. We remark that 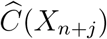 is valid, i.e., 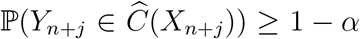, if all the data 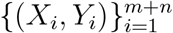 follow the same distribution. Therefore, CB ignores the distribution shift between labeled and unlabeled instances common in drug discovery tasks. Second, the probability is over all test samples instead of being averaged over the promising candidates. Hence, CB also ignores the selection effect.

The second baseline TU is the variant of TxConformal with *w*_*i*_ ≡ 1. That is, the selection set is the output of the Benjamini-Hochberg procedure applied at level *α* to 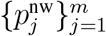, where 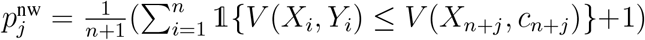. While this approach addresses the selection effect, it ignores the distribution shift between labeled and unlabeled data.

### A.7 Models and datasets

This subsection describes the AI models and datasets used in our experiments for Figures 2 and 3, together with the data splitting method to emulate realistic distribution shifts and performance evaluation strategies.

#### High-stability protein selection

For protein stability prediction, we use the pre-trained `ESM` protein language model^35^ within the TorchDrug PEER benchmark framework^77^ and finetune based on the task at hand. The model was trained on the protein stability dataset from Rocklin *et al*. (Science, 2017)^78^, which measures experimental stability changes across mutational variants. We calibrate on the pre-trained model’s validation set (proteins from found rounds of experimental design) and evaluate selection on the held-out test set (top candidate proteins with single mutations). We seek protein sequences whose true (unknown) stability falls in the top 20% of stability values in the training set (i.e., the cutoff *c* ∈ ℝ is defined as the 80-th percentile of stability values in the training set. We use the learned 1,280-dimensional `ESM` embeddings for the entropy balancing procedure. We then compute the false discovery rate (FDR) and power of selecting high-stability variants, where the results are averaged over *N* = 500 random subsamples of the calibration and test set within the original datasets (leaving the original distribution shift intact).

#### Gene perturbation selection

For selecting combinatorial gene perturbations, we use the `GEARS` model^41^, a graph neural network trained to predict cell fitness following genetic perturbations. We evaluate on the Norman *et al*. (2019) dataset of CRISPR-based combinatorial perturbations^40^, which systematically assays pairwise gene knockdowns in human cells. We adopt the split provided by GEARS, which partitions perturbation pairs uniformly into calibration and test sets, reflecting exploratory settings where combinations are sampled broadly without prior biological knowledge. The model is trained with a hidden size of 64, mean squared error loss, and batch sizes of 32 (training) and 128 (testing). We define “high-fitness” perturbations (the cutoff *c* ∈ ℝ) as those in the top 1% of true post-perturbation viability and construct selection sets accordingly. Unlike other tasks, no explicit distribution shift adjustment is needed, since perturbations are assigned randomly across calibration and test sets. We use the learned hidden representations from GEARS for entropy balancing in the conformal selection procedure. The procedures are repeated for *N* = 500 independent runs, where in each run the calibration and test sets are randomly split.

#### Enhancer sequence selection

For enhancer sequence selection, we use the pre-trained `Malinois` convolutional neural network (CNN)^46^. The model predicts enhancer activity in three cell lines, SK-N-SH, K562, and HepG2 and was trained on a dataset of 200 base pair enhancer sequences whose activity was measured via massively parallel reporter assay (MPRA). We calibrate on the pre-trained model’s validation set of enhancers from chromosome 13 and evaluate selection on chromosome 7, which was heldout from the model. We run selection three times, once for each cell line. In each cell line, we seek sequences in the top 10% of specific activity, as measured by activity in the target cell line minus the activity in the second most active cell line. We use 3,072 dimensional `Enformer`^49^ embeddings for the entropy balancing procedure. The procedures are repeated for *N* = 500 independent runs.

#### Generated promoter sequence selection

In our promoter experiment, we seek high activity sequences from promoter DNA sequences generated by an autoregressive model `RegLM`^52^. `RegLM` was trained to conditionally generate 80 base pair promoters at given activity levels in yeast in two media. As our candidate set, we generate 986 sequences using `RegLM` conditioned to have high activity in yeast in both media. As our scoring model, we use an `Enformer`^49^ model fine-tuned for `RegLM` for predicting yeast promoter activity in both media. We use the average of its predictions for both media to rank sequences. In lieu of experimental validation, we use a separate `Enformer` model trained on a non-overlapping set of yeast promoter data to label promoter activity. We calibrate on the test set of the prediction model and select sequences from the `RegLM` generated sequences. For entropy balancing embeddings, we use 512 dimensional embeddings from the 5th-to-last layer of the Polygraph^79^ yeast DNA sequence embedding model. We use top 10% label-model predicted activity as our selection cutoff. The procedures are repeated for *N* = 500 independent runs.

#### Clinical trial selection

For predicting trial outcome under temporal shift, we use `RelBench`’s relational trial-outcome task^80^ (`rel-trial-outcome`) built from the `rel-trial` dataset. We train a relational GNN (2 layers, 128 channels, sum aggregation) with `BCEWithLogitsLoss`, mini-batches of 512, and 10 epochs, incorporating text features via a `GloVe`-based text embedder. We adopt a time-based split in which earlier trials form the calibration set and later trials form the held-out test set, reflecting prospective deployment. To eliminate the impact of randomness, we repeat the procedure for *N* = 500 times, where in each replica the used calibration and test sets are a random subset of 75% of the original splits, which preserves the emulated distribution shift. Within each split, we designate “preferable outcome” trials as those meet a success criterion (binary label equals 1) and evaluate selection with TxConformal under temporal shift. We report empirical FDR and power on the held-out test period, averaged over *N* = 500 runs. For entropy balancing, we use the model’s hidden representations extracted from the NeighborLoader inference pass (dimension 128) on calibration and test trials.

#### CYP2D6 compound selection

For compound selection under scaffold shift, we use `AttentiveFP` (via `DeepPurpose`) as the predictive model^61^ and the `CYP2D6_Veith` dataset from the Thera-peutics Data Commons (TDC)^81^, a binary ADMET classification task indicating CYP2D6 inhibition. We perform scaffold splits using Bemis–Murcko scaffolds with an 80/10/10 train/validation/test partition on SMILES (`create_scaffold_split`), and then further split the training portion 90/10 into train/validation for model fitting, leaving the original 10% validation split as the calibration set and the held-out 10% as the test set. In this way, the training, calibration, and test sets contain compounds with distinct scaffolds. We train `AttentiveFP` for 50 epochs and extract 512-dimensional hidden embeddings (`get_hid_emb`) for the entropy-balancing procedure. Within each split, we target compounds in the top 10% of predicted inhibition probability as “high inhibition” and evaluate selection using TxConformal with a classification conformity score (binary cross-entropy loss). Conditional on the scaffold split, to reduce the randomness in evaluation, we randomly sample 75% of the original calibration and test sets, and use them as the calibration and test sets in the procedures. This is repeated for *N* = 500 independent runs, and we report false discovery rate (FDR) and power averaged over these replica.

### A.8 Methods for prospective deployment

#### Pre-experiment data processing

We began with the Enamine Hit Locator Library (HLL-460) with 460,160 molecules and filtered so the ensemble mean predicted activity was ≥ 0.1, resulting in 21,240 candidates. In order to ensure the novelty of our candidates, we further filtered the subset and preserve candidates that were ≤ 0.65 Tanimoto similarity to any molecule in the training and calibration data from Swanson et al. This left 21,206 remaining molecules. Finally, we used a procedure inspired by Sphere Exclusion Clustering^82^ to ensure the diversity of the final candidate set. Beginning with the molecule with the highest predicted activity, we repeatedly included the remaining molecule with the lowest Tanimoto similarity to any previously included molecule. We repeated this process until no molecules remained with ≤ 0.65 Tanimoto similarity. This resulted in a final candidate set of 14,978 molecules, which serves as the test set in TxConformal.

#### Pre-experiment model development

We here detail the model development process. We start with eight candidate prediction models, which are deep neural networks trained with eight embeddings `AttentiveFP`, `CNN`, `GCN`, `GIN_AttrMask`, `GIN ContextPred`, `Morgan`, `NeuralFP`, `rdkit 2d` using the DeepPurpose library^62^. To ensure model robustness, we emulate realistic shifts and examine the performance of TxConformal using each of the models. Specifically, we design three data splitting strategies to produce the synthetic training/calibration/test data from the Swanson et al. data, including (i) `Random_All`: random split, which introduces no distribution shift, (ii) `Scaffold_All`: scaffold split, so that the training, calibration, and test data all have distinct scaffolds, and (iii) `Calib_Test_Scaffold`: a mixture of random and scaffold split, where we first randomly split the entire set to obtain the training fold, and then scaffold split the remaining data to obtain the calibration and test fold. We then use the synthetic training fold to train each of the eight property prediction model using DeepPurpose^62^. For each property prediction model, TxConformal uses its hidden embeddings and predicted values to estimate the weights and construct p-values. We then run TxConformal to select test samples at various FDR target levels, and evaluate the realized FDR on the test fold using the known labels. The entire procedures are repeated for *N* = 100 independent runs, and we evaluate the empirical FDR as the average of fraction of false positives across those runs. We find that two GIN models `GIN_AttrMask` and `GIN_ContextPred` lead to robust empirical FDR control and superior selection power. The final prediction model is set to be the ensemble of the two GIN models (the model prediction is the average of the two GIN model predictions), which yields robust empirical FDR control with the ensemble model under the three data splitting regimes (SI Figure 5) in this semi-synthetic evaluation.

#### Pre-experiment analysis with T**x****C****onformal**

We then apply TxConformal before experimental validation, where the calibration data is from Swanson et al.^65^ and the test data is the filtered candidate set described above from the Enamine HLL library. Following the final evaluated approach, TxConformal takes (the empirical distribution of) the mean predictions and the last-layer embeddings of the two models to learn the weights. TxConformal then produces p-values for each test compound following the method in Section A.3, and ranks the compounds in ascending order of the p-values. The top-*K* candidates are selected for synthesis based on the budget *K* = 100. After selection, we use the method for scenario 4 in Figure 3 to produce the point estimate for the number of false positives by *p*_[*K*]_ · *m*, where *p*_[*K*]_ is the largest conformal p-value among selected compounds, and *m* = 14, 978 is the total size of the candidate pool. This produces a point estimate of 80.30 false positives. We then apply the normal approximation formula and obtain a 90% confidence interval of [70.00, 94.60]. For comparison, we apply the unweighted variant of TxConformal, which produces a point estimate of 46.80 false positives. A normal approximation leads to a 90% confidence interval of [32.50, 61.11] for the number of false positives. The conformal baseline does not provide estimation of the error rates, and is not directly applicable here. However, mimicking a scientist who select compounds whose prediction sets at marginal coverage level 1 − *α*, it takes *α* = 0.012 to select these top-100 compounds (this is an approximation, since the weights slightly change the ranking of the compounds). As such, the scientist who used the conformal baseline would think the error rate is 0.012.

#### Antibacterial activity prospective experiment

Acinetobacter baumannii ATCC 17978 was grown overnight in 3 mL of LB medium at 37°C with shaking. The bacteria were diluted 1:10,000 in fresh LB and added to 96-well flat-bottom polystyrene plates (Corning) containing compound or DMSO alone (0.5%, solvent control), for a final volume of 100 µL and a final test compound concentration of 50 µM. The plates were incubated statically at 37°C under humidified conditions for 18 h before OD600 was measured using a BioTek Synergy Neo2 MicroPlate Reader (Agilent). To calculate the level of inhibition, the background absorbance (medium alone, average of at least 8 wells) was first subtracted from all wells. Percent growth was then calculated as the ratio of each treated well to the average of the DMSO-treated control wells (8 wells).

#### Post-experiment analysis

After receiving the experimental results, we compute the fraction of false discoveries of 0.91 (91 out of 100 molecules have an inhibition of *<* 20% averaged over two replicates). We evaluate the selected molecules using three criteria: activity, novelty, and diversity. We analyze the selected active candidates via pairwise Taniomoto distance among them and their novelty scores defined as the minimal Tanimoto distance from training and calibration data.

## Supplementary Notes

## B Supplementary results for experiments

## C Theoretical results

In this part, we derive asymptotic theory for our p-values and error-controlling procedures. These results generalize the finite-sample results under knowledge of the sampling process such as exchangeability and known distribution shift^1, 2^.

*Proof of Theorem 3*. The first conclusion is a re-statement of the conformal selection framework^1^. We now consider the general setting with score *V* (*x, y, c*). Note that *V* (*x, y*; *c*) is non-decreasing in *y*, and thus 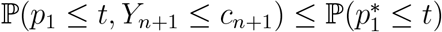, where

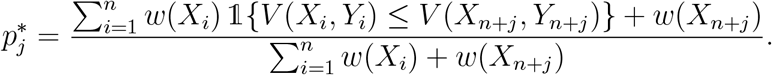

We now verify 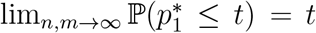 under the given conditions. By the theory of entropy balancing^83^, the solution to (1) with *δ* = 0 is of the form 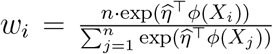 for some data-dependent 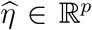. Under condition (i), according to^83^,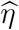 converges to a fixed constant *η* ∈ ℝ^*p*^ and thus by the Cauchy-Schwarz inequality,

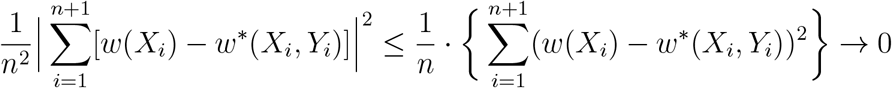

in probability since 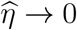 as *n, m* → ∞. On the other hand, similarly,

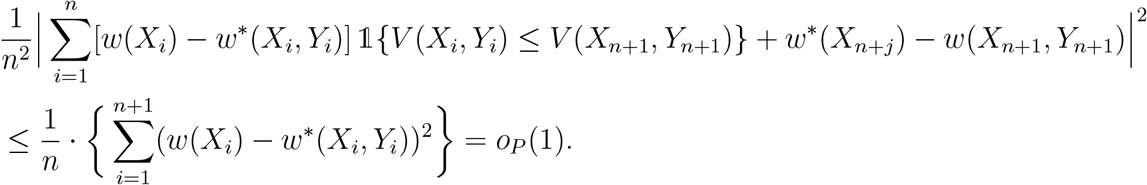

These together imply 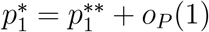, where

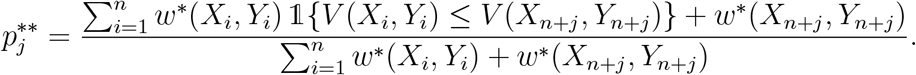

Using the theory of conformal inference under weighted exchangeability^84^, since *w*^∗^(*x, y*) is the true density ratio, we know 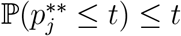 for any *t* ∈ [0, 1]. Therefore, we have 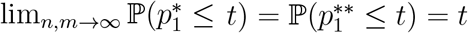, which completes the proof under condition (i).

Now suppose condition (ii) holds, in which case 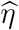 still converges to a fixed constant *η* but the true density ratio is not necessarily proportional to exp(*η*^⊤^*ϕ*(*x*)). We define the convergence limit of *w*(*X*_*i*_)’s by the function 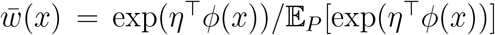. Then, similar to previous arguments as well as the uniform law of large numbers, we can show 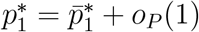, where

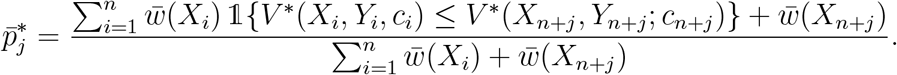

Under condition (ii), conditional on 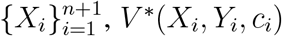’s are i.i.d. random variables, and the weights 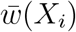 are fixed. Thus, by the theory of conformal inference under exchangeability^15^, conditional on the unordered set of 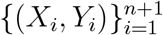, and letting [1], [2], …, [*n* + 1] be an ordering of (1, …, *n* + 1) such that *v*_[*k*]_ is the *k*-th order statistic of 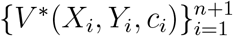, we know that the random variable 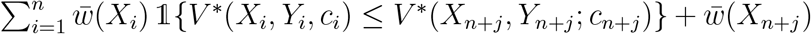 follows a uniform distribution on (the finite set)

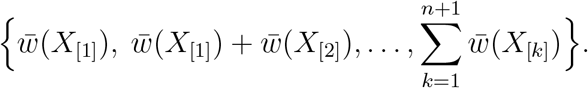

This implies that 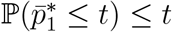 for any *t* ∈ [0, 1], which further implies 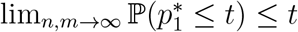, completing the proof under condition (ii).

*Proof of Theorem 4*. The first conclusion under *P* = *Q* is a re-statement of the conformal selection framework^1^. We now consider the general estimated case. By the definition of the Benjamini-Hochberg procedure, the selection set is equivalently^85^

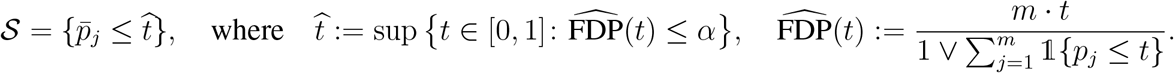

Thus, the false discovery proportion obeys

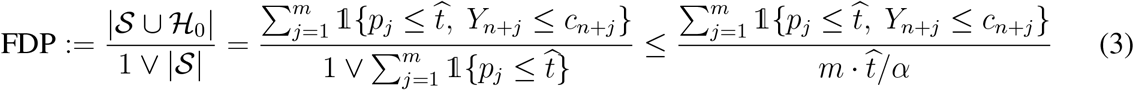

by the definition of 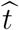. We will show that the upper bound will be asymptotically controlled.

By the theory of entropy balancing^83^, the solution to (1) with *δ* = 0 is of the form 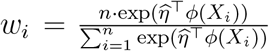 for some data-dependent 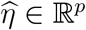, and 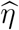 converges to a fixed *η* ∈ ℝ^*p*^ as *n, m* → ∞. Now let 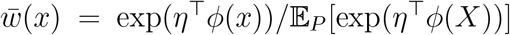 be the limiting function. Therefore, by the Cauchy-Schwarz inequality,

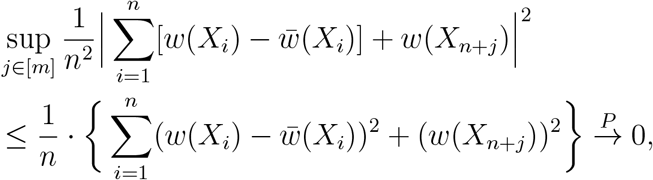

where 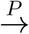 represents convergence in probability, and

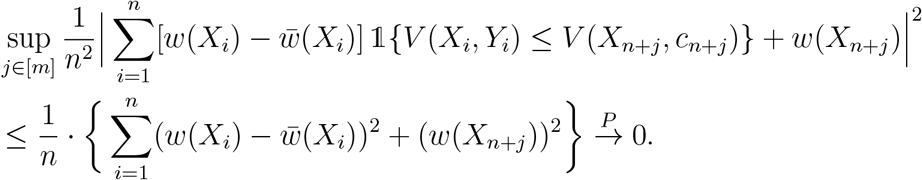

By the law of large numbers, we have 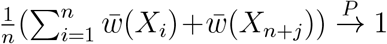. In addition, by the uniform law of large numbers for cumulative distribution functions, we have

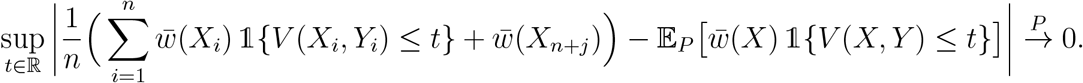

These facts imply that 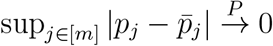, where we define

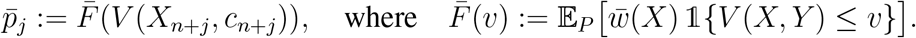

Therefore,

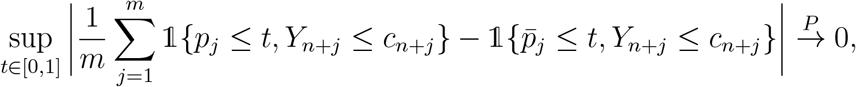

which further implies (by the uniform law of large numbers)

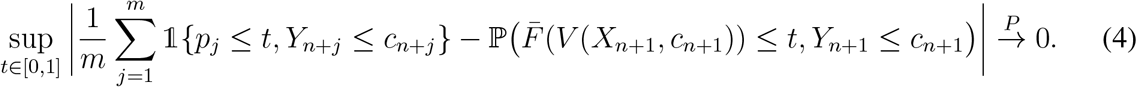

With similar arguments, we can show that

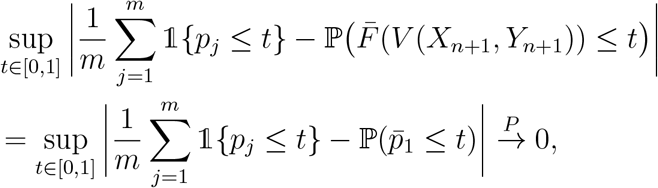

which further implies that for any *δ >* 0,

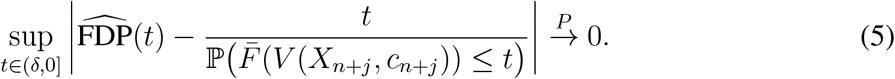

Define 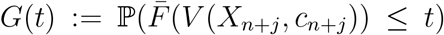 and 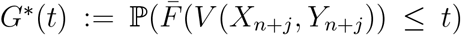. By the definition of 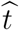, under regularity conditions on 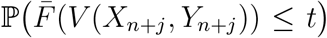 similar to the common ones in the literature^2, 85^, 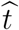 is lower bounded away from zero and converges to a fixed constant:

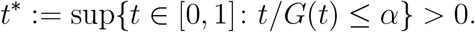

Due to the monotonicity of *V* (*x, y*) in *y*,

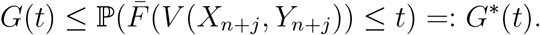

Under condition (i), we know 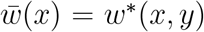, hence 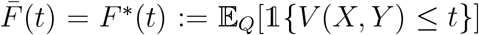. This means *G*^∗^(*t*) = *t*, since 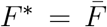 is the c.d.f. of the random variable *V* (*X*_*n*+*j*_, *Y*_*n*+*j*_). Combined with (3), (4) and (5) and the fact that 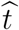 is bounded away from zero, this means

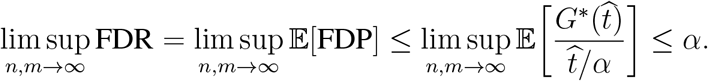

Now suppose 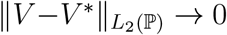. We can show that 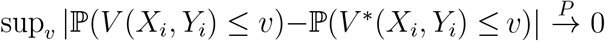. Let 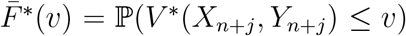 and *G*^∗∗^(*t*) = ℙ(*V* ^∗^(*X*^*n*+*j*^, *Y*^*n*+*j*^) ≤ *t*). We know that 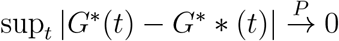.

Under condition (ii), since the conditional distribution of *V* ^∗^(*X*_*i*_, *Y*_*i*_; *c*_*i*_) is invariant given *X*_*i*_, we have 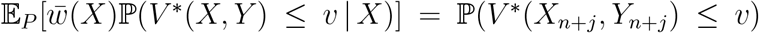 for any function 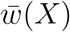, which further implies *G*^∗∗^(*t*) = *t*. Similar arguments as those under condition (i) then yield lim sup_*n,m*→∞_ FDR ≤ *α* under condition (ii).

Finally, taking an asymptotic argument for the balancing condition imply that 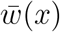 always obeys 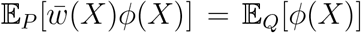, Let 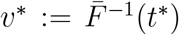. Under condition (iii), we know 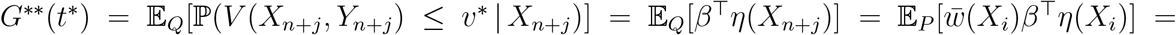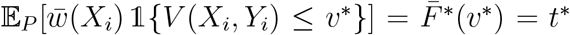. Similar arguments as in the preceding two cases then yield lim sup_*n,m*→∞_ FDR ≤ lim sup_*n,m*→∞_ *G*(*t*^∗^)*/*[*t*^∗^*/α*] ≤ *α* under condition (iii).

*Proof of Theorem 5*. By the definition of 𝒮, it is equivalently 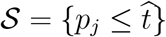, where

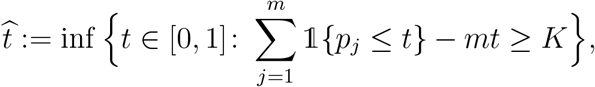

and we define inf(∅) = 1. Thus, when 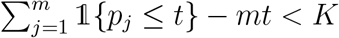 for all *t* ∈ [0, 1], 𝒮 = [*m*] and |𝒮 ∩ ℋ_1_| = |ℋ_1_|; otherwise, letting *H*_0_ = {*j*: *Y*_*n*+*j*_ ≤ *c*_*n*+*j*_}, the definition of 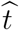 implies

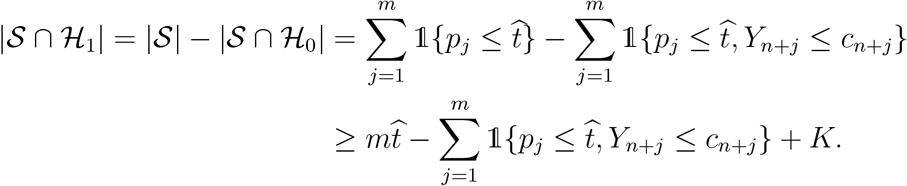

Following similar arguments as in the beginning of the proof of Theorem 4, we have

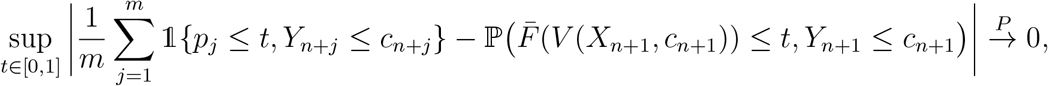

where 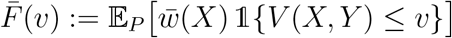, and

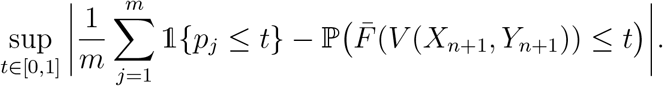

This implies 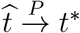, where we define the fixed constant 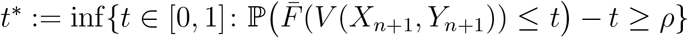, and recall that inf(∅) = 1 and *K/m* → *ρ*. (When 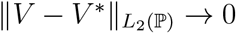, *V* is replaced by *V* ^∗^ in the definition of *t*^∗^.)

Under conditions (i), similar to the proof of Theorem 4, 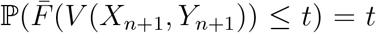 for any *t* ∈ [0, 1], which implies 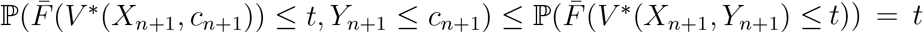. This further implies lim inf_*n,m*→∞_ |𝒮 ∩ ℋ_1_|*/m* ≥ lim inf_*n,m*→∞_ min{|ℋ_1_|*/m, K/m*}, which yields the desired conclusion as *K/m* → *ρ*.

Under condition (ii), similar to the proof of Theorem 4, we can show 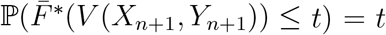 for any *t* ∈ [0, 1] where 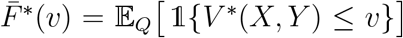, which also leads to the desired conclusion following the arguments under condition (i).

Finally, under condition (iii), using the fact that 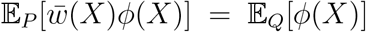, we have 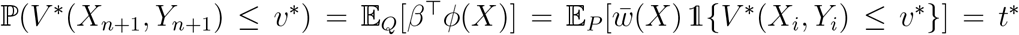, which also leads to the desired conclusion.

*Proof of Theorem 6*. The selection set is equivalently 𝒮 = {*p*_*j*_ ≤ *K/m*}. Therefore,

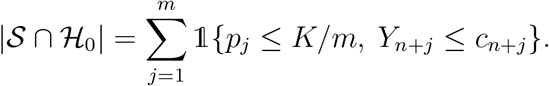

With the same arguments as in the beginning of the proof of Theorem 4, we know

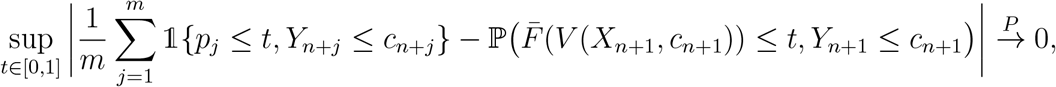

where we recall that 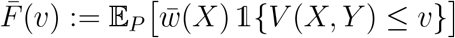.

Therefore, we know that 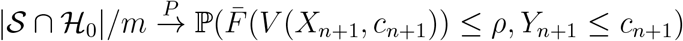, which is upper bounded by 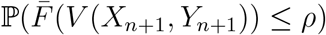 due to the monotonicity of *V* (*x, y*).

Similar to the proof of Theorem 4, condition (i) implies 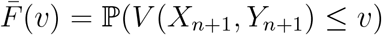 for any *v*, and hence 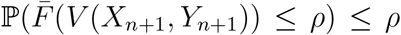. Therefore, lim sup_*n,m*→∞_ |𝒮 ∩ ℋ_0_|*/m* ≤ *ρ*, which is equivalent to lim sup_*n,m*→∞_ |𝒮 ∩ ℋ_0_|*/K* ≤ 1 since *K/m* → *ρ*.

When 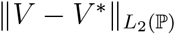, we can similarly show that

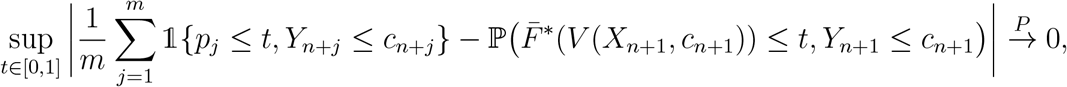

where 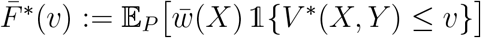. Thus, under condition (ii), we also have 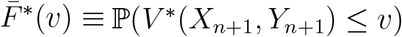 and thus lim sup_*n,m*→∞_ |𝒮 ∩ ℋ_0_|*/K* ≤ 1.

Finally, under condition (iii), the balancing condition and the linear assumption imply 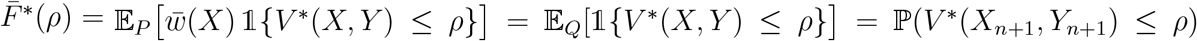, which yields the desired conclusion lim sup_*n,m*→∞_ |𝒮 ∩ ℋ_0_|*/K* ≤ 1.

*Proof of Theorem 7*. With the score specified in this part, when *M* is a sufficiently large constant so that *M >* 2 sup_*x*_ *f* (*x*) and *M >* 2*c*, we know that *V* (*X*_*i*_, *Y*_*i*_) ≤ *V* (*X*_*n*+*j*_, *c*) only if *Y*_*i*_ ≤ *c*. Therefore, the p-values are equivalently

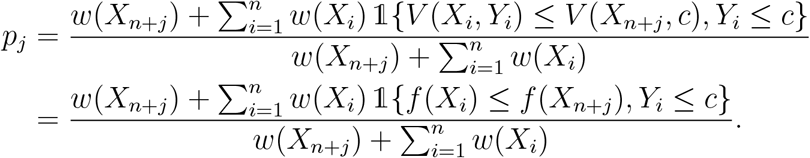

With similar arguments as in the proof of Theorem 4, we can show that 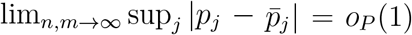 where 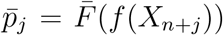, and we define 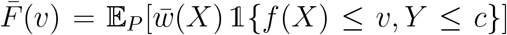.

Combined with the uniform law of large numbers, this implies

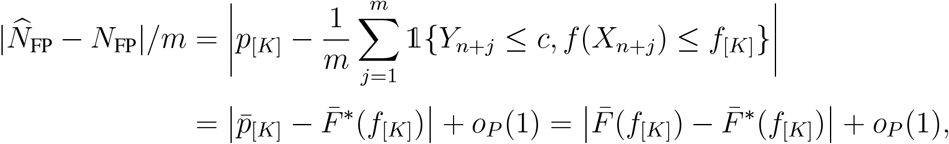

where 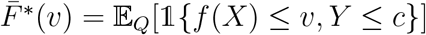.

Both conditions (i) and (ii) imply 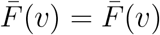 hence 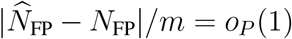, which leads to the desired result as *K/m* → *ρ*. On the other hand, with *K/m* → *ρ* we can show that 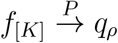 where *q*_*ρ*_ is the *q*-th quantile of *f* (*X*) under *X* ∼ ℚ. Thus 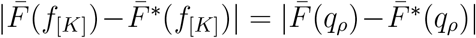. Using the balancing condition for 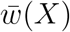, under condition (iii), we also have 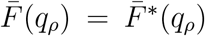 thereby leading to the desired result.

